# Massive experimental quantification of amyloid nucleation allows interpretable deep learning of protein aggregation

**DOI:** 10.1101/2024.07.13.603366

**Authors:** Mike Thompson, Mariano Martín, Trinidad Sanmartín Olmo, Chandana Rajesh, Peter K. Koo, Benedetta Bolognesi, Ben Lehner

**Author notes:** (B.B) and (B.L.).

## Abstract

Protein aggregation is a pathological hallmark of more than fifty human diseases and a major problem for biotechnology. Methods have been proposed to predict aggregation from sequence, but these have been trained and evaluated on small and biased experimental datasets. Here we directly address this data shortage by experimentally quantifying the amyloid nucleation of >100,000 protein sequences. This unprecedented dataset reveals the limited performance of existing computational methods and allows us to train CANYA, a convolution-attention hybrid neural network that accurately predicts amyloid nucleation from sequence. We adapt genomic neural network interpretability analyses to reveal CANYA’s decision-making process and learned grammar. Our results illustrate the power of massive experimental analysis of random sequence-spaces and provide an interpretable and robust neural network model to predict amyloid nucleation.

## Introduction

Specific insoluble protein aggregates in the form of amyloid fibrils characterize more than fifty clinical conditions affecting more than half a billion people (Fig. 1A)^1^. These include common neurodegenerative disorders and the most frequent forms of dementia. Nonetheless, amyloids are present in all kingdoms of life and can have functional roles, including in humans^2^. Protein aggregation is also a major problem in biotechnology, for example in the production of enzymes, antibodies and other protein therapeutics^3^. The importance of amyloids across biological functions and diseases has spurred massive research efforts, yet the determinants and mechanisms of their formation remain quite poorly understood^4,5^.

**Figure 1.**
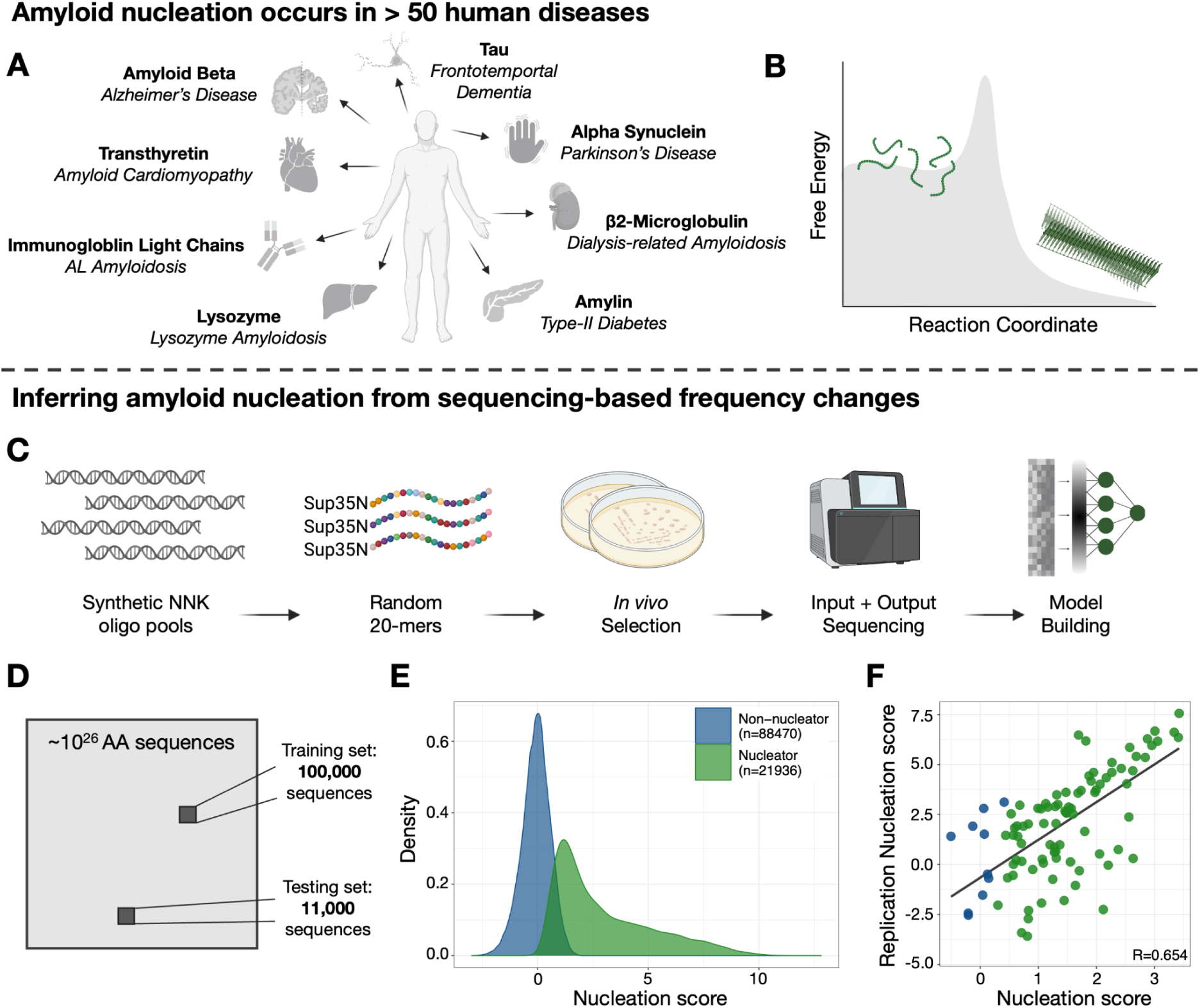
Quantifying the nucleation of >100,000 random peptides. (A) Examples of amyloids in human diseases. (B) The amyloid state is thermodynamically favorable, but requires overcoming a kinetic barrier. (C) Experimental design. (D) While we explore over 110,000 sequences, our dataset is a tiny sample of the possible sequence space. (E) The assayed nucleation scores of sequences labeled “Nucleators” and “Non-nucleators” in our experiment. (F) An example of a follow-up replication experiment using a synthesized library (NNK3; see Supplementary Fig. 1 for others; Supplementary Data File 3).

Recent advances in cryogenic electron microscopy have allowed the atomic structures of many mature amyloid fibrils to be determined^6^. Amyloids share a cross-β structure wherein hydrogen-bonded β-strands are perpendicularly stacked along the fibril axis, creating β-sheets that face each other and are parallel to the fibril axis^4,7,8^. Amongst humans, amyloid fibrils typically have hydrophobic cores, for which hydrophobicity and β-strand propensity form the basis of many computational methods to predict amyloid propensity from sequence^9–15^. However, other amyloids, for example yeast prions, have very different sequence composition, hinting at a richer diversity of amyloid-forming sequences^16,17^.

In contrast to the remarkable advances in the structural characterization of mature fibrils, the process of amyloid formation—how soluble proteins overcome a free energy barrier to nucleate fibrils (Fig.1B)—is much less understood. Time-resolved structure determination has been used to study the *in vitro* assembly of amyloids, revealing a striking diversity of intermediate structures appearing and disappearing as fibrillation proceeds^18,19^. However, how this process initiates and why it only occurs for some sequences under physiological conditions remains unclear. Mature amyloid fibrils are very stable and are likely to be the thermodynamically favored state at high protein concentration for many proteins^20,21^. There is, however, a very high energy barrier to amyloid nucleation for most proteins i.e. the process is under kinetic control^21^. The kinetic control of amyloid nucleation is, therefore, the key problem to understand: what are the sequence-level determinants that cause some peptides to nucleate amyloid formation on timescales relevant to biology?

We believe that our ability to understand and predict amyloid formation is currently data-limited. To date, computational methods to predict aggregation have been trained and benchmarked on very small and biased experimental datasets, making it unlikely they have learnt representations of aggregation that generalize across the diversity of sequence space^12–15,22,23^. For example, even for 20 amino acid sequences there are 20^20^ (>10^26^) different sequences of 20 amino acids. Such a large sequence space is unlikely to have been accurately modeled by methods trained on tens or a few hundred sequences.

To directly address this data gap we have developed a massively parallel selection assay that allows the aggregation of thousands of different proteins to be tested and quantified in a single experiment^24,25^. This has allowed us to quantify the nucleation kinetics for all possible substitutions, insertions and deletions in the amyloid beta peptide that aggregates as a hallmark of Alzheimer’s disease. The resulting measurements agree very well with *in vitro* nucleation kinetic rate constants^24,25^. However, these datasets are limited to testing the effects of small changes to a single sequence, hindering utility for general-purpose model-building.

Here we apply this approach at a much larger scale and quantify the nucleation of >100,000 peptides with completely random sequences. We use the resulting massive dataset to evaluate existing aggregation prediction methods and find they are only moderately predictive. We therefore use the data to train CANYA, a convolution-attention hybrid neural network. This fast model dramatically outperforms existing predictors of protein aggregation when tested on >10,000 additional sequences, demonstrating the power of massive experimental sequence-space exploration. Subsequent post-hoc explainable AI (xAI) analyses provide mechanistic insights into CANYA’s decision-making process and learned grammar. CANYA provides a robust and interpretable neural network model for understanding and predicting amyloid-forming proteins. More generally, our results provide a very large and well-calibrated dataset to train and evaluate models beyond CANYA and they demonstrate the utility of massive experimental analysis of random protein sequence-spaces.

## Results

### Massively parallel quantification of amyloid nucleation kinetics for >100,000 sequences

To better understand the sequence determinants of amyloid nucleation kinetics, we used an in-cell selection assay to quantify the rate of nucleation of more than a hundred thousand peptides with fully random sequences. We generated four libraries (NNK1-4) of random 20 AA peptides using NNK degenerate codons (where N = A/C/G/T and K = G/T) and expressing them as fusions to the nucleation domain of Sup35 (Sup35N), a yeast prion-forming protein that allows fitness-based selection for amyloid nucleation (Fig. 1C)^24–26^. Briefly, fusion sequences that nucleate amyloids sequester Sup35 resulting in translational readthrough of a premature stop codon in the *ade1* gene so that cells containing those sequences become able to survive in medium lacking adenine. Enrichment or depletion of each sequence after selection can be quantified by deep sequencing, with enrichment scores linearly related to the log of *in vitro* amyloid nucleation rates^24,25^.

Each library was selected independently and sequencing was used to quantify the relative enrichment (‘nucleation score’) for each genotype in the library. Sequences in the first three experiments made up our training and testing sets (NNK1-3, ∼111,000; Fig. 1D; Supplementary Data Files 1 and 2), corresponding to about a 1/10^17^ fraction of the possible sequence space (20^20^), while sequences from the fourth experiment (NNK4, ∼7,000) were used as a held-out validation data set. After data processing and quality control, the vast majority of sequences had a nucleation score of 0. Consequently, we classified sequences with a nucleation score significantly greater than 0 (one-sided Z-test, FDR adjusted p-value <=0.05) as nucleators (n=21,936), and all other sequences as non-nucleators (n=88,470) (Fig. 1E). Importantly, these nucleation scores are reproducible, as measured by an additional selection experiment on a designed library (replication library) re-quantifying the nucleation of 400 sequences sampled across all four libraries (Pearson correlation range 0.506-0.797, Fig. 1F, Supplementary Fig. 1).

### Nucleating sequences span a large sequence-space and are poorly predicted by existing computational methods

After classifying sequences as nucleators and non-nucleators, we sought to characterize each class through amino acid composition (Fig. 2A), physicochemical properties (Fig. 2B), and current amyloid prediction tools (Fig. 2C).

**Figure 2.**
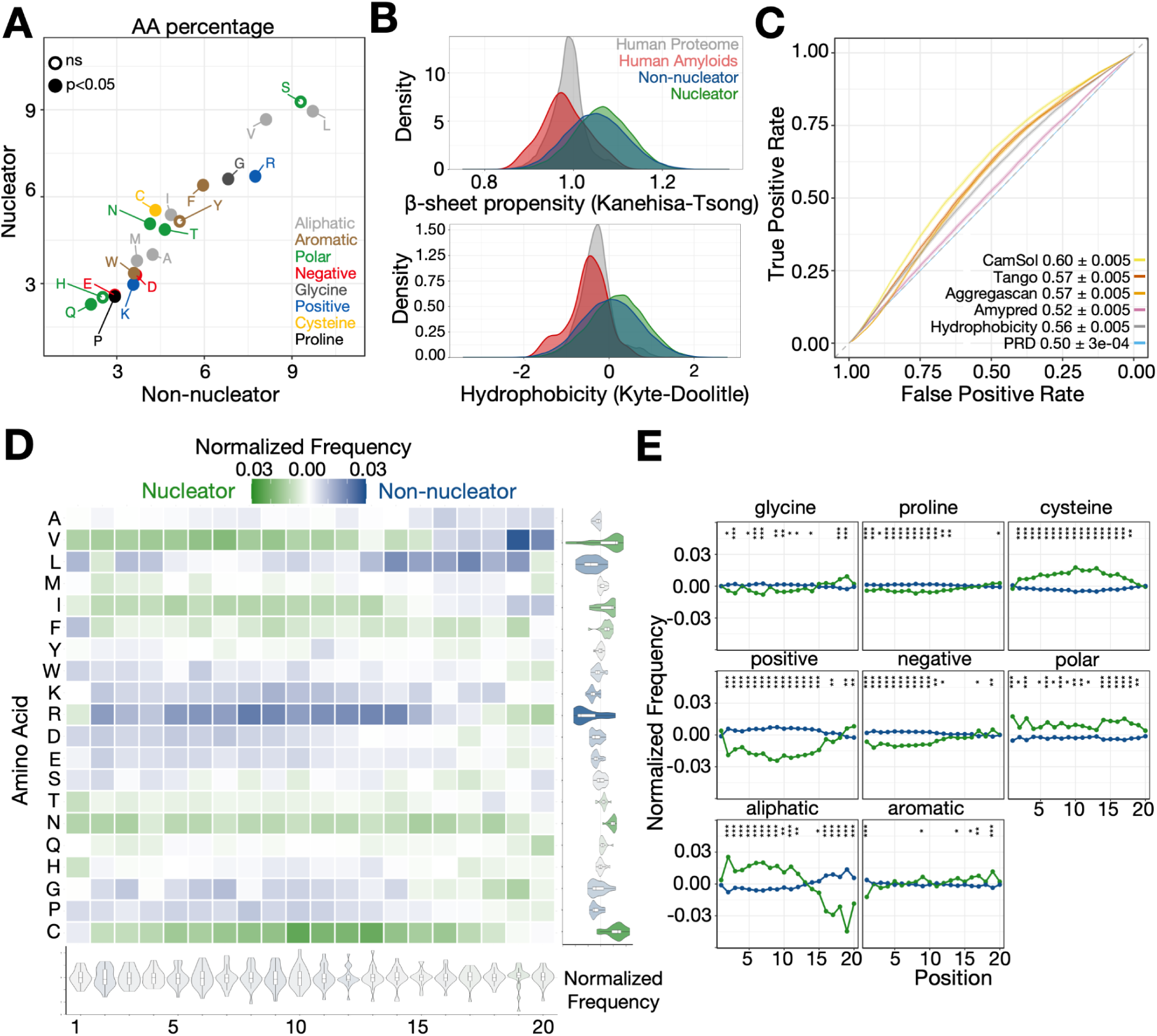
Nucleation is poorly predicted by existing models and subtly related to amino acid composition. (A) The percent composition of residues grouped by their physicochemical properties in nucleators and non-nucleators. (B) The hydrophobicity and beta-sheet propensity of assayed sequences relative to known human amyloids (Supplementary Table 2) and the human proteome. (C) The predictive power (AUC ± 95% CI) of previous amyloid predictors on the random sequences. (D, E) The position-specific differences in amino acid frequencies across nucleating and non-nucleating sequences. Asterisks indicate marginal p-value (chi-square test) lower than 0.05 “*”; lower than 0.01 “**”; lower than 0.001 “***”.

First, we examined the differences in amino acid frequency between nucleating and non-nucleating sequences. Differences in frequencies were generally modest, however we observed statistically significant differences owing to the large sample size of our data. When looking at composition independent of position, nucleators had higher frequencies of cysteine (difference in frequency 0.012, p<2e-16), asparagine (0.009, p<2e-16), and isoleucine (0.005, p<2e-16), and lower frequencies of arginine (-0.010, p<2e-16), leucine (-0.008, p<2e-16), and lysine (-0.006, p<2e-16; Fig. 2A, See Supplementary Table 1 for full differences). Moreover, both nucleators and non-nucleators covered the beta-sheet propensity and hydrophobicity spaces of the human proteome and known amyloid sequences, and nucleators had slightly higher values of both than non-nucleators on average (difference in means of hydrophobicity=0.130, beta-sheet propensity=0.012, both two-way t-test p-values < 2e-16; Fig. 2B). Considering position-specific composition, differences were again modest, ranging from a difference in frequency from -0.06 to 0.03 (Fig. 2D). Subsequently, we grouped amino acids by their physicochemical properties to check for more broad, position-specific differences between the two sequence classes (Fig. 2E). Toward the N-terminus of the random sequence (i.e., closer to Sup35N), nucleators were significantly enriched (chi-squared test) for aliphatic residues (min p-value=1.54e-13, position 2 difference=0.033), and significantly depleted for positive (min p-value=1.57e-25, position 9 difference=-0.032) and negative residues (min. p-value=3.14e-11, position 2 difference=-0.016). The differences in charge waned toward the C-terminus (min. p-value above position 15=1.03e-3, position 20 charged difference=0.011), however, and frequency differences in aliphatic residues changed such that nucleators were significantly *depleted* for aliphatic residues relative to non-nucleators (min. p-value=5.77e-39, position 19 difference=-0.058). Several groupings showed other position-sensitive differences, such as an enrichment of aromatic residues toward the C-terminus in nucleators (min. p-value=5.09e-6, position 19 difference=0.015), an enrichment of varying strength for polar residues in nucleators (p-value= 5.57e-8 position 1 difference=0.023, p-value=9.41e-7 position 17 difference=0.020), and the enrichment of cysteines away from the ends of the random construct (min. p-value=1.11e-28, position 10 difference=0.023).

Despite statistical significance, we highlight that differences in sequence space are subtle. In other words, the collection of slight variation in amino acid frequencies offers minimal insight or definitive conclusions around the overall properties or characteristics determining nucleation in our experiment. To attempt to elucidate characteristics that separate the sequence classes and consequently learn important axes of variability, we turned to dimensionality reduction techniques. In addition to manually examining differences within the first several dimensions, we also used the scores in lower-dimensional space as features in a logistic multiple regression task to distinguish nucleators from non-nucleators. Using principal components analysis (PCA), we observed no clear separation between nucleators and non-nucleators whether we used amino acid composition alone (cumulative variance explained from the top 10 PCs = 54.7%, Area Under ROC curve (AUC) using all 10 PC scores=0.601, 95% Confidence Interval (CI)=[0.596, 0.607], Supplementary Fig. 2), or maintained positionality of the amino acids when fitting the model (cumulative variance explained from the top 10 PCs = 3.1%, AUC=0.564, 95% CI=[0.559, 0.570], Supplementary Fig. 2). This modest separation between classes of sequences was consistent even when using non-linear embedding techniques (first 10 UMAPs AUC=0.584, 95% CI [0.578, 0.589]), or adding amino acid propensities to the dimensionality reduction tools (first 10 PCs AUC=0.614, 95% CI [0.608, 0.619]; Supplementary Fig. 2).

As dimensionality reduction methods were unable to distinguish the classes of sequences, we next explored whether separation is possible using existing amyloid predictors. Beyond hydrophobicity indices, several of these methods include structural information^27^ or model biophysical mechanisms^12^, potentially enabling them to capture more complex features of nucleation. We applied several state-of-the-art amyloid prediction algorithms to our data and found that the methods either failed to generalize to our data or had only modest predictive power (Fig. 2C, CamSol, highest AUC=0.598, 95% CI [0.593, 0.603]). We posit that, since many of these tools have been trained on very small sets of known amyloids or moderate numbers of short hexamer sequences, their applicability to our experimental data may be limited. To understand where the methods underperformed, we examined the scores from the highest performing methods (CamSol^13^ and TANGO^12^) and found that non-nucleating sequences with a high-predicted nucleation score had higher hydrophobicity (two-sided t-test p-value<2e-16) than all other non-nucleating sequences (Supplementary Table 3). We also found that low-predicted nucleators had higher presence of positive (two-sided t-test p-value<2e-16) and negative (p-value<2e-16) residues than all other nucleators (Supplementary Table 4).

### CANYA: a hybrid neural-network that accurately predicts amyloid nucleation

Given previous approaches failed to accurately predict nucleation status within our dataset, we built our own model to capture the sequence-nucleation score landscape. Concretely, we developed a hybrid neural network which we term CANYA, or Convolution Attention Network for amYloid Aggregation. Though a neural network may seem inherently less interpretable than simpler models, as we explain below, the architecture of CANYA is not only simple, but also biologically motivated. CANYA builds off the observation that known amyloids are composed of interacting short sequences, such as stacked beta sheets, and treats this information as an inductive bias for the model—first the sequences are passed through a convolutional layer which discovers ‘motifs’, then these motifs are passed through an attention layer to learn positional effects of motifs and to encourage these motifs to interact with each other (Fig.3 A). Moreover, we set the filter lengths of the convolutional layer based on the distribution of secondary structure lengths in 80 known amyloid fibril structures (WALTZ-DB^28^, Supplementary Fig. 4). Though—to our knowledge—this class of models is new to proteins, convolution-attention hybrid models have been used in genomics and found to serve as a sound inductive bias for discovering motifs and their interactions^29,30^.

**Figure 3.**
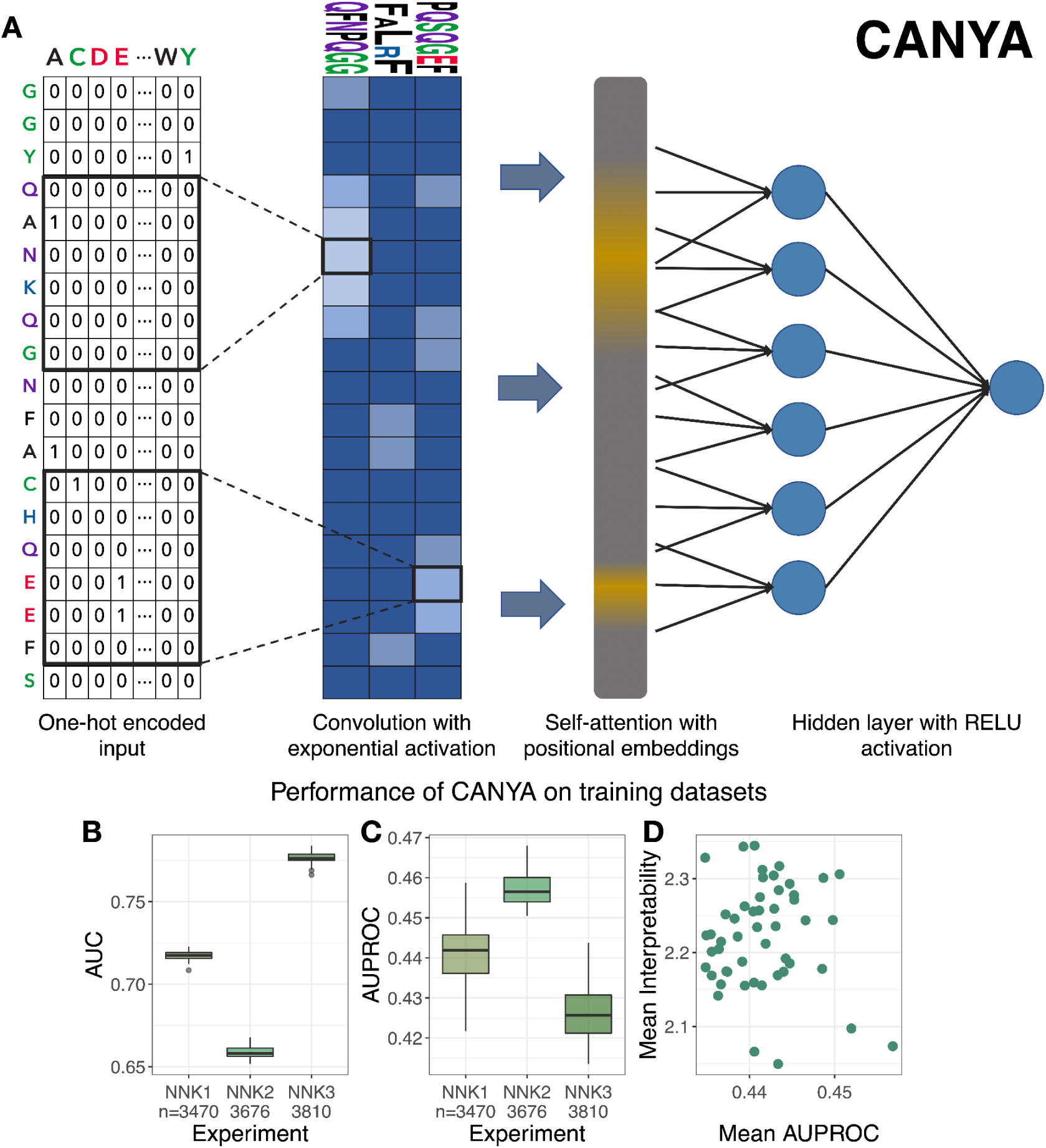
Convolution-Attention Network of amYloid Aggregation (CANYA). (A) CANYA is a 3-layer neural network with 65,491 parameters. The model contains 100 filters, a single attention head with key-length 6, a dense layer with 64 nodes, and finally a sigmoid output layer. (B-D) Evaluation metrics across the top 50 performing (of 100) model fits of CANYA. (B) The area under receiver operating characteristic curve (AUC) for held-out testing sequences. (C) The area under precision recall curve (AUPROC) for held-out testing sequences. (D) The interpretability score (KL divergence; Methods) calculated on all held-out test sequences plotted against the mean AUPROC across experiments. See Supplementary Fig. 3 for results on all 100 model fits.

We trained CANYA 100 times on over 100,000 synthetic sequences and their respective nucleation status to learn the sequence-nucleation landscape. Unlike massive, computationally intensive neural networks, CANYA comprises only three layers (spanning 65,491 parameters) and requires less than an hour to train on a basic, modern CPU. Despite this simplicity, and having only observed a small fraction of the possible sequence space, CANYA substantially improved the prediction of nucleation status of held-out test sequences (average AUC=0.710, 0.650, 0.769 across NNK experiments 1-3 respectively, Fig. 3B-C) over previous methods (max AUC CamSol, NNK1=0.617, NNK2=0.537, NNK3=0.673). We also note that the predictive accuracy of CANYA was significantly higher than simpler linear models trained on the same dataset with amino acid composition or counts alone (Supplementary Fig. 5).

To understand the differences in performance across methods, we examined the sequence scores between the next best performing method (CamSol) and CANYA. We found that the largest discrepancies for non-nucleating sequences occurred in hydrophobic sequences with tryptophans, and in cysteine-or asparagine-rich sequences with few aliphatic residues in the case of nucleating sequences (Supplementary Tables 3 and 4). Our results not only highlight the utility of exploring a vast sequence space, but also suggest that CANYA is able to contextualize physicochemical properties within sequences (e.g., among hydrophobic sequences, CANYA adjusts its score in the presence of bulky, or disruptive residues).

Crucially, we developed CANYA with the goal of interpreting the grammar of nucleation rather than maximizing predictive power. We accordingly scored each trained instance of CANYA using a recently developed interpretability metric to select a model amenable to uncovering this learned grammar^31^. Briefly, this metric examines the enrichment of motifs utilized when training the model and compares them to the set of all equal-length (k=3) kmers in the training sequences (Methods). Strong enrichment (i.e., divergence from the background training sequences) indicates a model may yield clearer resolution in downstream interpretability analyses. Though the area under the precision-recall curve (AUPROC) of test sequences was more consistent than AUC across experiments (average AUPROC NNK1=0.434, NNK2=0.452, NNK3=0.415 ; Fig. 3C), we did not find a correlation between predictive performance and this interpretability metric (correlation of average AUPROC and interpretability score r=-0.059, p-value=0.6847, Fig. 3D). We therefore chose the trained model with the highest interpretability score, conditional on the fact that it scored better than the median-performant model (of 100 training runs; Methods).

### Evaluation on >7,000 additional sequences

To further evaluate the performance of CANYA and to compare it to that of previous methods we quantified the nucleation of an additional ∼7,000 random sequences (Fig. 4A). The sequence spaces spanned by the training and these test sequences are effectively independent (∼10^5^ and ∼10^3^ samples from a >10^22^ sequence landscape). CANYA remained highly accurate on the 7,000 unseen sequences (AUC CANYA=0.809, 95% CI [0.798, 0.821; Fig. 4B, PROC in Supplementary Fig. 6). Moreover CANYA substantially outperforms all tested previous methods^12–15,23,32^. The next best performing method was Aggrescan (AUC=0.707 95% CI [0.694, 0.719]), followed by TANGO (AUC=0.680 [0.667, 0.693]) and CamSol (AUC=0.679 [0.665, 0.693]). Neither AmyPred nor PLAAC produced significantly accurate predictors on the validation dataset, which may be indicative of over-fitting on their respective training datasets—we used a simple hydrophobicity score as a baseline predictor, which scored AUC=0.593 95% CI [0.579, 0.607]).

**Figure 4.**
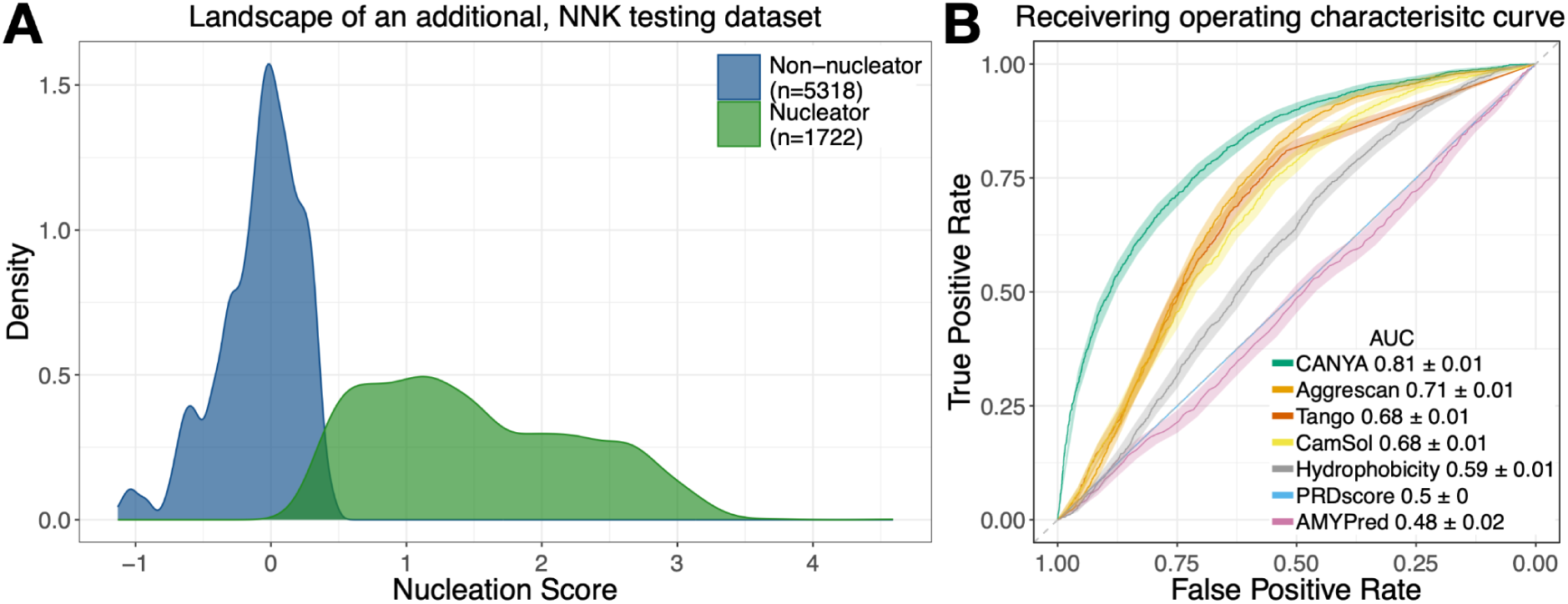
An additional experiment of >7,000 random sequences. (A) The nucleation rates over an additional validation set of 7,040 random sequences. (B) The predictive performance (AUC ± 95% CI) of CANYA and previous methods on the additional dataset.

### CANYA predicts known amyloids and aggregating sequences

After establishing that CANYA can accurately predict the experimental nucleation status from primary sequence, we sought to understand whether the nucleation function learned by CANYA is applicable to longer sequences and different contexts. We first considered 1,400 hexapeptides from WALTZ-DB, the largest previous dataset of amyloidogenic and non-amyloidogenic sequences^28^. Strikingly, however, on these six-AA peptides no method significantly outperformed hydrophobicity for classifying aggregating from non-aggregating sequences (AUC=0.813 05% CI [0.791, 0.836]) (Fig. 5). The hydrophobicity distributions of amyloid and non-amyloid hexamers in WALTZ-DB are indeed very distinct (Supplementary Table 5), suggesting biases in this dataset or that hydrophobicity dominates the aggregation potential of such very short peptides. This cautions against the use of such short sequences for model training and evaluation.

**Figure 5.**
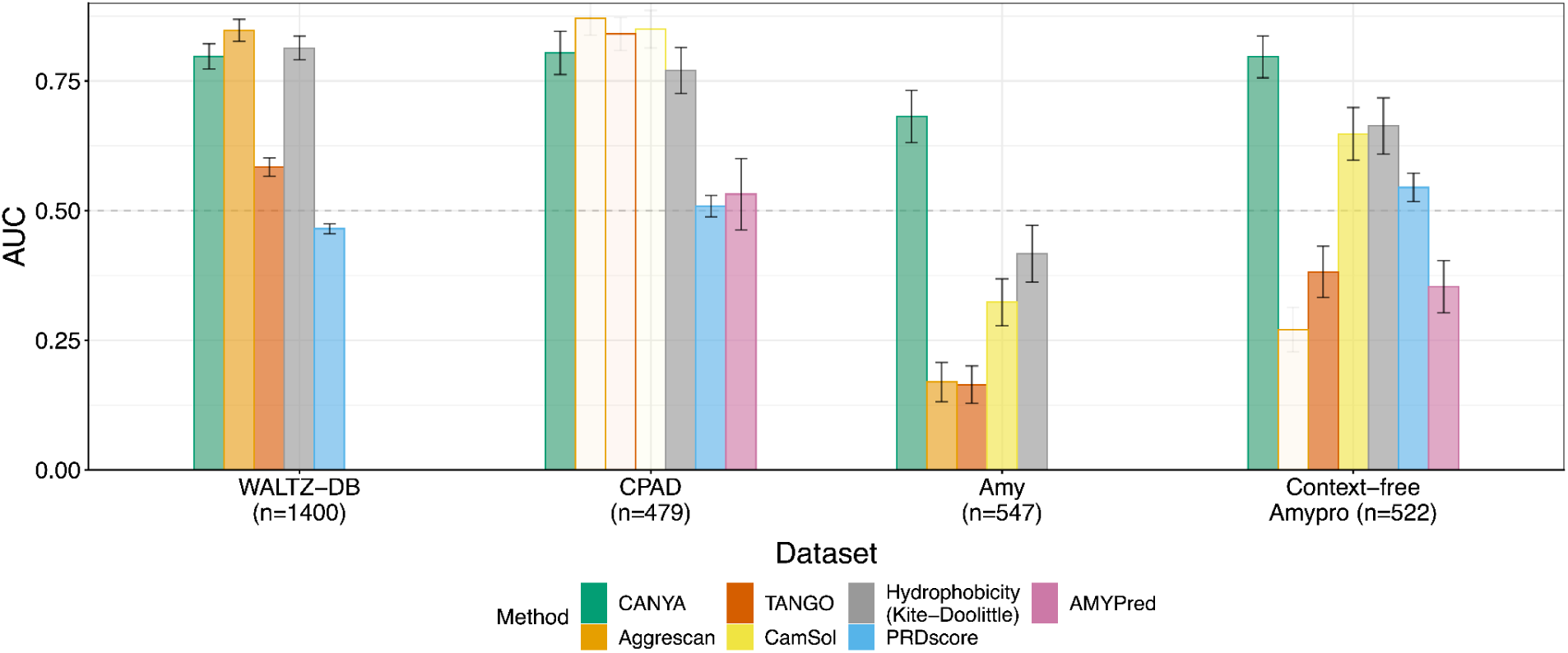
Stable performance of CANYA across diverse prediction tasks. The AUC of CANYA and previous methods across several external datasets. Low-opacity bars represent cases in which the method used data from the testing dataset for training and thus are not valid out-of-sample evaluations. See text for additional descriptions of datasets (Methods, Supplementary Table 5) as well as performance reported as area-under precision-recall curve (AUPROC; Supplementary Fig. 6).

We next considered the Curated Protein Aggregation Database (CPAD). Though CPAD contains over 2,000 sequences, we limited our evaluation here to the 479 sequences with length >10 AA (median length 16 (Q1 length=10 and Q3 length=22), comprising 304 amyloid-forming sequences and 175 non-aggregating sequences (Supp Table 5). It is important to note that several of the previous methods (including TANGO, CamSol, and Aggrescan) were directly trained on sequences within CPAD, violating the ability to evaluate their out-of-sample predictive performance on this dataset. Despite this, CANYA performed similarly as well as these methods on CPAD (AUC=0.804, 95% CI [0.762, 0.845], AUPROC=0.855, 95% CI [0.817, 0.890]) (Fig. 5).

We also evaluated performance on the Amy dataset^33^ which contains a set of much longer sequences, including 382 non-amyloid sequences (median length=708 [Q1=344.5, Q3=1375.5]) gathered from UniProt and 165 amyloid sequences (median length=162 [Q1=77, Q3=443]) from the AmyPro database^34^. Strikingly, CANYA was the only predictor with both statistically significant AUC and PROC in this task (AUC=0.681 95% CI [0.631, 0.731], Fig. 5, AUPROC 0.495, 95% CI [0.428, 0.568]). The poor performance of hydrophobicity and previous methods suggests the importance of features beyond sequence composition in determining amyloid propensity in longer protein sequences.

Finally, we evaluated whether each method could identify amyloidogenic regions of each protein in the AmyPro dataset. Specifically, we evaluated whether methods were capable of distinguishing an amyloidogenic region from a non-amyloidogenic region in the absence of any contextual region (Methods), for which we term the task “Context-Free AmyPro.” CANYA significantly out-performed all previous approaches (AUC=0.796, 95% CI [0.756, 0.837], Fig. 5), and was the only method to significantly out-perform hydrophobicity on this task (AUC=0.663, 95% CI [0.609, 0.717]).

In summary, CANYA’s performance is state of the art and consistent across diverse prediction tasks and protein sizes. Moreover, CANYA is significantly more accurate than simpler linear models that are trained over the same NNK training set (Supplementary Fig. 5)

### CANYA learns physicochemical nucleation motifs

We next performed a series of interpretability analyses to understand how CANYA assigns its nucleation score and to elucidate difficult-to-see patterns that differentiate the nucleators and non-nucleators in the training data.

First, to visualize physicochemical ‘motifs’ learned by the model, we constructed position-weight-matrices (PWMs) using kmers that activated a given filter at least 75% of the maximum-activating kmer (Methods). We selected a filter length of 3, as this is the mode length of secondary structures in structurally resolved amyloids (Methods; Supplementary Fig. 4). Motifs showed clear physicochemical preferences (Fig. 6). For example, many motifs capture blocks of hydrophobicity (clusters 1 and 2) or charge (clusters 6 and 8). Some motifs showed heterogeneity, or position-preferential effects, such as polar or charged residues being surrounded by hydrophobic (clusters 4 and 5) or aromatic residues (clusters 7, 9 and 10; Fig. 6).

**Figure 6.**
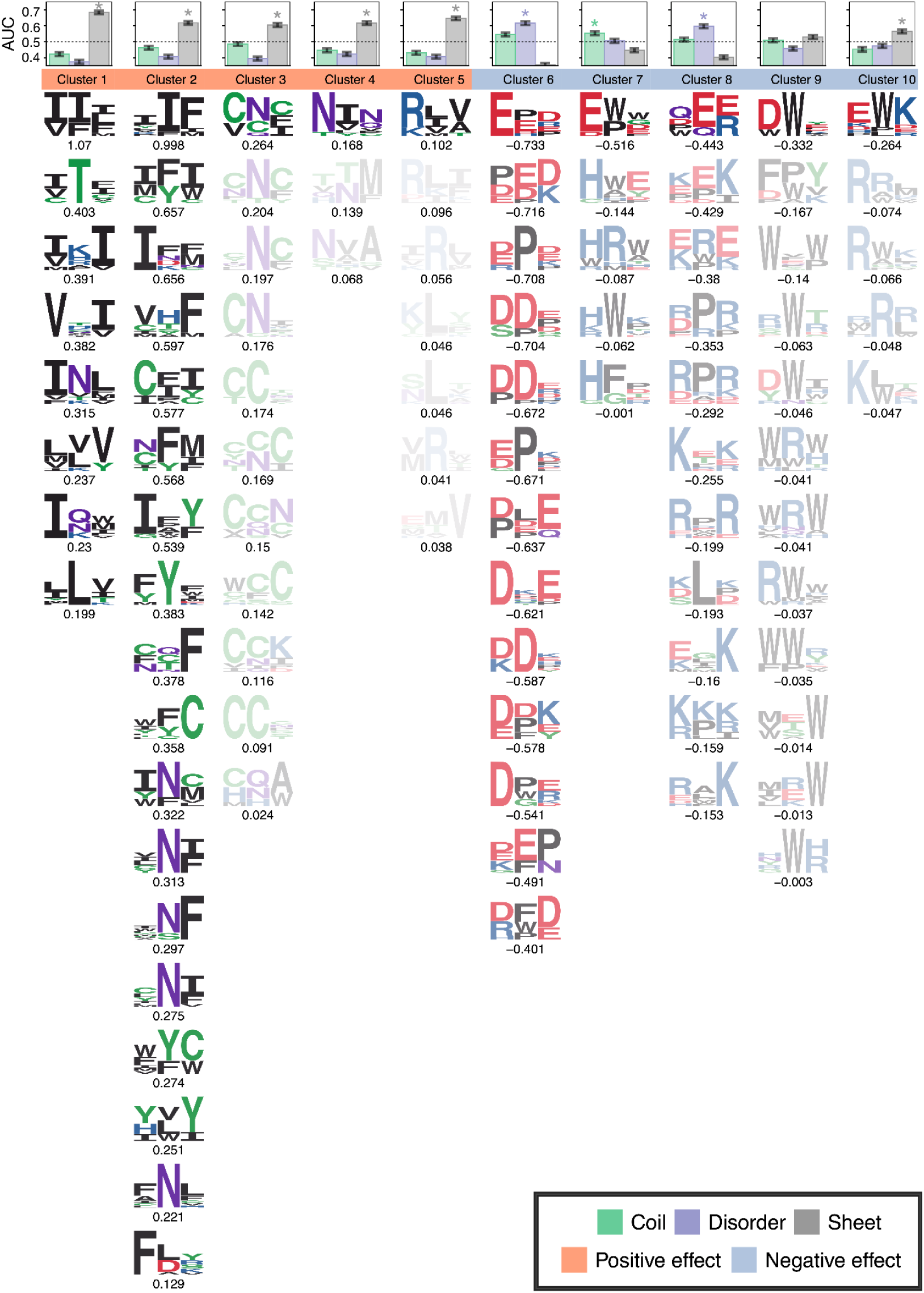
CANYA discovers physicochemical nucleation motifs. The motifs discovered by CANYA, clustered by their physicochemical properties and GIA effect sizes, then sorted based on their effect size magnitude. Translucency represents the ratio of cluster effect size compared to the strongest cluster (Methods). The enrichment (in AUC) of motif-cluster presence in secondary structures of resolved amyloids in Uniprot (Methods; Supplementary Fig. 7). The dashed lines represent an AUC of 0.50 and asterisks represent structures for which the enrichment was significantly higher than both 0.50 and the second most-enriched structure.

We next turned to a post-hoc interpretability method named Global Importance Analysis (GIA) to learn the effect of each motif^35^. Briefly, GIA learns effect sizes by embedding a motif of interest in a set of background sequences, then comparing the difference in the models’ predicted nucleation propensity between these background sequences with and without the embedded motif (Fig. 7A). The effects learned by CANYA recapitulated previously known amyloid biology—hydrophobic motifs strongly increased a given sequence’s propensity to nucleate, and charged, proline-containing motifs lowered sequences’ propensity to nucleate (Fig. 6)^36–38^. Motifs containing residues enriched in yeast prions (Q/N) also increased amyloid propensity (weaker motifs of clusters 1, 2, and 3, stronger motifs of cluster 4), as did motifs enriched in cysteine (cluster 3) or aromatic residues (cluster 2; Fig. 6). Interestingly, CANYA could also uncover motifs for which specific residues had effect sizes in both directions. For example, tryptophan-containing motifs led to a negative effect when the tryptophan was surrounded by charged residues (clusters 7, 9, 10; Fig. 6), or a positive effect in the context of hydrophobic, polar, or other aromatics (clusters 1, 2, 3; Fig. 6). Notably, CANYA also found a set of motifs enriched in hydrophobicity with a positively charged residue (cluster 5, Fig. 6), further suggesting the model captures previously uncharted areas of the amyloid sequence space.

**Figure 7.**
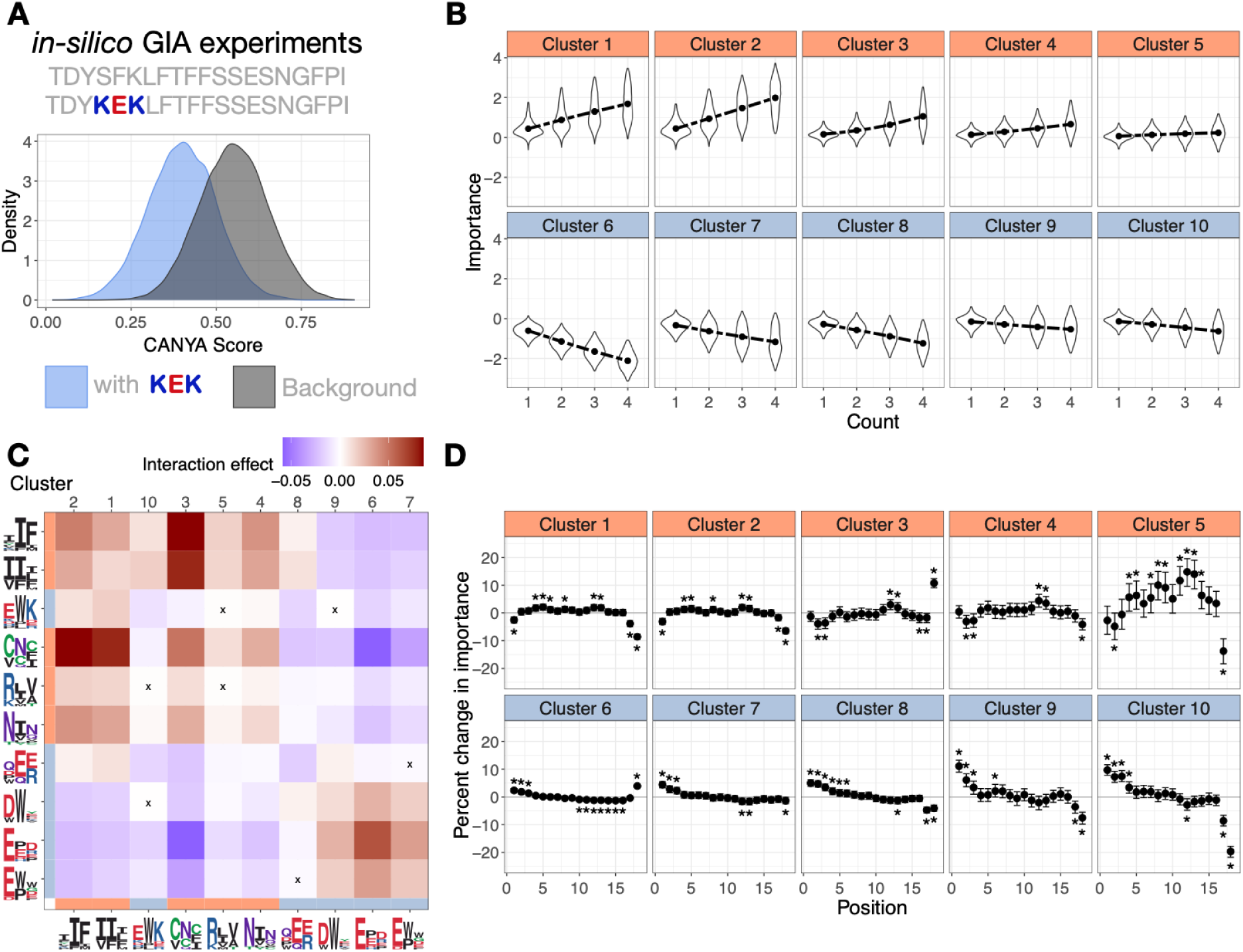
*in-silico* experiments reveal CANYA’s learned nucleation grammar. (A) An example of an experiment using GIA, an explainability tool to extract importance (effect sizes) of features in a model. Briefly, model predictions for a background set of sequences are compared to predictions on the same set of sequences with a feature (motif) embedded in them. (B) The distribution of effects from adding 1-4 copies of a cluster-motif to sequences. Points represent importance. (C) Interaction importance from adding motifs from two clusters to sequences. Warmer colors indicate higher CANYA score than from marginally adding the motifs (and their effects) separately to sequences, whereas cooler colors represent a CANYA score lower than expected from adding marginal motif effects. “X” indicates effects that were not significantly different from 0. (D) The position-dependence of motif effects. Plotted is the percent change of a position-specific effect relative to the motif’s global, position-averaged effect. Stars represent a significantly non-zero percent change in effect.

We next clustered the motifs using their AA similarity to generate a more concise representation of what the model has learned and to reduce the dimensionality of *in silico* analyses to extract further information learned by CANYA. To do so, we first generated BLOSUM scores (which capture a similarity of amino acids based on evolutionary divergence) for each motif, then performed affinity clustering on the BLOSUM scores to derive a candidate set of clusters (Methods)^39^. We verified that this approach results in a sound set of clusters by re-running GIA using the clusters as the feature of interest and confirming that the learned effect size for a cluster was consistent with the motifs of which it is composed (Methods; Supplementary Table 6). We were left with 10 clusters on which to perform downstream *in silico* experiments, effectively reducing the number of experiments by at minimum one order of magnitude (from 100 filters).

### Physicochemical motif activation in known amyloid structures

We examined whether the motif clusters discovered by CANYA showed propensity for secondary structures in known amyloid fibril structures (from the Structural analysis of Amyloid Polymorphs (StAmP) database^40^). We included in our comparison full-length resolved structures of amyloid fibrils for 114 PDB entries comprising amyloid structures of 23 proteins (Supplementary Table 7). Here, we used the activation energy of a cluster across positions to predict whether or not the corresponding position was in a beta-strand, other structured region (coil), or unresolved (disordered, see Methods). The AUC from this task serves as a metric of whether high activation (high matching score) of a motif is associated with a specific structural element. Clusters with high hydrophobicity and positive effect size were most strongly associated with activating in beta strands (Fig. 6; Supplementary Fig. 8, max AUC=0.683, 95% CI [0.672, 0.693], cluster 1), whereas the strongest enrichment amongst negative-importance clusters was observed in disordered regions (max AUC=0.617, 95% CI [0.605, 0.628], cluster 6). Interestingly, negative-importance clusters containing tryptophan showed varying enrichments in secondary structures (Fig. 6; Supplementary Fig 8). Clusters with tryptophans near histidines were moderately enriched in coils (cluster 7; AUC=0.553, 95% CI [0.541, 0.565]), tryptophans adjacent to positive charges showed moderate enrichment for strands (cluster 10; AUC=0.566, 95% CI [0.554, 0.577]), and tryptophans near other aromatics (cluster 9) showed no significant enrichment for any structure.

### Motif position-dependence and interactions

Treating the motif clusters as input for GIA, we performed an additional set of experiments to evaluate whether CANYA has learned positional information of motif effects and whether motif effects are additive (Fig. 7A).

To learn positional information for each cluster of motifs, we ran an experiment in which we calculated the GIA effect of the cluster at every position of the construct, and compared it to the global, position-averaged effect of the cluster. These comparisons revealed that CANYA was also able to learn position-relevant information across each cluster of motifs (Fig. 7D). Generally, the positive-effect clusters showed diminished effects at the ends of the construct and stronger effects at the center (Fig. 7D). The range of percent change was most drastic for cluster 5, potentially due to the presence of a charged residue (%-change in effect from 14.81% 95% CI [9.59, 19.59] to -39.81 %, 95% CI [-43.76, -36.02]; Fig. 7D). Clusters 1, 2, and 4 followed similar trends, however the changes were much more modest (highest percent change position 12, cluster 4=4.42%, 95% CI [2.05, 6.73], lowest percent change position 18, cluster 1=-8.58%, 95% CI [-9.53, -7.63]). Cluster 3, which is marked by high presence of cysteines, followed a similar trend except that its effect significantly increased in the last position (% change=10.76, 95% CI [9.11, 12.37]).

Conversely, the negative-importance clusters all had strengthened effects toward the N-terminus, where the peptides are fused to Sup35N,, and all but the proline-rich cluster (cluster 6) had diminished effects toward the C terminus (Fig. 7D). This may be due to the fact that cluster 6 was the most negatively charged cluster, consistent with negative charges in the C-terminus of some amyloid-forming peptides reducing fibril formation^41^.

Tryptophan-containing motifs (clusters 7, 9, and 10) typically had increased effects closer to the N-terminus (greatest % change cluster 9, position 1=11.14% 95% CI [8.96, 13.28]) and dampened effects at the C-terminus (greatest % change cluster 10, position 18=-19.67 [-21.74, -17.80]). Like cluster 5, cluster 10 contains positively-charged and hydrophobic residues and had its most dampened effect at position 18. However, cluster 10’s effect size is in the opposite direction, likely due to the presence of tryptophan. To learn whether the effects of motifs were additive, we ran an experiment where we embedded motifs in a cluster in non-overlapping positions between 1 and 4 times. Simple additive effects explained nearly all of the variance observed in model predictions (range of R^2^ between multiplicity and importance = [0.971, 0.999]; Fig. 7B). However, some clusters showed evidence of heteroskedasticity in their importance values, which may indicate minor epistatic or background-specific grammars. Accordingly, we used GIA to perform an experiment similar to the one determining additivity of motifs, however we focused on the case in which there are only two motifs, and the embedded motifs are selected from different clusters (Methods). This enables us to learn how interactions between clusters affect nucleation scores. Every cluster showed at least 8 statistically significant interactions (p-value of paired, two-way t-test <0.05 /(10*10 tests); Fig. 7C; Methods), suggesting the importance of modeling sequence context in the prediction of nucleation status. Nonetheless, cluster interaction effects were modest (ranging from -0.058 to 0.085) compared to cluster main effects (-0.608 to 0.448). Almost all clusters exhibited a self-enhancing effect in which their interaction importance was significantly higher than the importance from additively combining each marginal effect (maximum importance cluster 6, 0.063). This was not the case for the mixed-charge disorder cluster (cluster 8; importance -0.012 p-value<2e-16) interacting with itself, nor the negative-importance charged, hydrophobic cluster (cluster 10; importance -0.011 p-value<2e-16). Interestingly, the hydrophobic and aromatic positive-importance clusters (clusters 1 and 2, respectively), showed positive interaction effects with the mixed-charge disorder cluster (cluster 8), suggesting that the interaction with disordered regions (like cluster 8) could facilitate amyloid nucleation by hydrophobic regions (like clusters 1 and 2)^42^. On the contrary, disorder clusters (clusters 6 and 8) showed negative interactions with the cysteine and asparagine positive clusters (clusters 3 and 4; FIg 7C).

## Discussion

Amyloid protein aggregation is a hallmark of many human diseases and a major problem in biotechnology. However, relatively few protein sequences are known to nucleate amyloids under physiological conditions, and this shortage of data likely limits our ability to understand, predict, engineer, and prevent the formation of amyloid fibrils.

Here we have directly addressed this data shortage by quantifying the nucleation of amyloids at an unprecedented scale (100,000 random sequences) and used the data to evaluate the performance of existing computational models. Finding the performance of these methods to be limited, we then used the data to train CANYA, a fast and interpretable deep learning model of amyloid nucleation. Evaluation on an additional independent 7,000 sequences confirmed the performance of CANYA but not other methods on predicting nucleation from sequence.

Using random sequences allowed us to test the nucleation of sequences very different to the small number of known amyloids and to provide a principled evaluation of existing amyloid predictors^12–15,23^ using both our own and existing datasets^23,28,34,43^, serving as a guideline for the community. The excellent performance of CANYA and its consistency across evaluation tasks suggests CANYA does in fact learn an accurate approximation of the sequence-nucleation landscape, despite only training on random, synthetic peptides. The performance of previous methods compared to that of hydrophobicity scales suggests that the use of limited dataset sizes and short peptides has limited the amount of additional nucleation-relevant information these approaches could learn. This underscores the importance of using longer sequences and high-throughput assays to profile previously unexplored regions of the sequence-nucleation landscape.

CANYA has an inherently interpretable model whose architecture is inspired by biology. We also adapted state-of-the-art explainable AI (xAI) techniques from genomic neural networks to the protein space^30,31,35,44^. This not only reveals insight into the decision-making process of our model, but also illustrates how xAI techniques developed for genomic neural networks can provide intelligible information from neural networks that model protein function.

Interpretability analyses identified ‘physicochemical motifs’ that underlie CANYA’s decision making process, including nucleation promoting motifs enriched in beta strands of known amyloid structures and nucleation preventing motifs enriched in disordered regions of known amyloids. The effects of these physicochemical motifs combined mostly additively, with only subtle motif-motif interactions, suggesting a modest role for long-range epistasis or context-specificity in the process of amyloid nucleation. However, the physicochemical motifs did have position-specific effects, and these warrant additional investigation in future experimental work.

A potential limitation of our strategy is that we only tested the nucleation of sequences of 20 amino acids and in one particular experimental context. There likely remains additional predictive power to be harvested by experimentally testing at scale and modeling longer sequences and consequently longer-range interactions. Nonetheless, we found through several evaluations that the information learned by modeling the length-20 constructs from our experimental assay can offer accurate predictions of nucleation status across a wide range of protein lengths and contexts.

An additional consideration is CANYA’s architecture. We limited our neural network architecture to a relatively simple class of models as our focus was on interpretability. Recent literature suggests that leveraging protein embeddings—in lieu of one-hot encoding sequences—may boost our predictive power^45–52^, though such an approach will likely pose difficulties when performing post-hoc xAI experiments as done here^53^. Further, our model comprises a modest 65,000 parameters and leverages sparsity despite having over 100,000 sequences on which to learn. Many models of protein structure employ much more complex architectures, with both substantially larger numbers of layers and parameters^45,50,52,54–56^. Future investigations may build off of the work presented here by generating longer sequences, or exploring more complex architectures.

The pairing of massive scale experimental data generation using random sequences with interpretable models has led to insights into genomic regulatory functions^57^. However, to the best of our knowledge, it has been little utilized in the space of proteins to probe mechanisms beyond short motifs. We believe the approach deserves wider adoption, whenever sequences are functional at sufficient frequencies to allow their identification in practical library sizes. Systematic large datasets such as the one presented here can be re-used to train and evaluate additional models, and the predictions and outputs of these models can loop back into additional large-scale experimental explorations of sequence space.

## Data Availability

All datasets generated from this study are provided under Gene Expression Omnibus (GEO) accession number GSE268261.

External datasets can be found under their respective repositories, which we list here:

AmyPred https://pmlabstack.pythonanywhere.com/dataset_AMYPredFRL

AmyPro http://www.amypro.net/

CPAD https://web.iitm.ac.in/bioinfo2/cpad2/index.html

WALTZ-DB http://waltzdb.switchlab.org/sequences.

## Code availability

CANYA is open-source, free to use, and available at the following link https://github.com/lehner-lab/canya.

## Author contributions

BB, BL, and MT conceived the project. MT and MM performed computational analyses with input from CR, PKK, BB and BL. TSO performed all experiments. MT conceived and wrote the model with input from CR, PKK, BB and BL. MT, MM, BB and BL wrote the manuscript, with input from all authors.

## Acknowledgements

This work was funded by the La Caixa Research Foundation project ‘DeepAmyloids’ (LCF/PR/HR21/52410004). Work in the lab of B.L. was also funded by an European Research Council (ERC) Advanced (883742) grant, the Spanish Ministry of Science and Innovation (LCF/PR/HR21/52410004, EMBL Partnership, Severo Ochoa Centre of Excellence), the Bettencourt Schueller Foundation, the AXA Research Fund, Agència de Gestió d’Ajuts Universitaris i de Recerca (AGAUR, 2017 SGR 1322), the CERCA Program/Generalitat de Catalunya and Wellcome (Grant reference: 220540/Z/20/A, ‘Wellcome Sanger Institute Quinquennial Review 2021-2026’). Work in the lab of BB was funded by the Spanish Ministry of Science, Innovation and Universities (PID2021-127761OB-I00, RYC2020-028861-I funded by MCIN/AEI/ 10.13039/501100011033, “ERDF A way of making Europe” and “ESF Investing in your future”) and by the European Union (ERC Consolidator, Glam-MAP, 101125484). MT was funded by EMBO Fellowship ALTF 266-2023. P.K. and C.R. were funded by NIH grants R01HG012131 and R01GM149921. We thank Andre J Faure for performing preliminary analyses and helpful suggestions.

## Methods

### Plasmid library construction

Libraries of random sequences (NNK1-4) were synthesized by Integrated DNA Technologies (IDT) as ultramers of 20 NNK codons (60 nucleotides). A library containing 400 sequences selected from the previous four random libraries was synthesized as an oligopool by IDT for validation and replication (Supplementary Data Files 3 and 4). In both cases, sequences were flanked by constant regions of 25 nt upstream and 21 nt downstream for cloning. The NNK ultramers and the replication oligo pool were extended in a 1-cycle PCR (Q5 high-fidelity DNA polymerase, NEB) with primers TSO_2 and TSO_65 (Supplementary Data File 5). The resulting products were treated with 2µl/tube of ExoSAP (ExoSAP-IT, Applied Biosystems) for 30 minutes at 37 °C and 20 minutes at 80 °C and purified through a MinElute column (Qiagen). In parallel, the PCUP1-Sup35N plasmid was linearized by PCR (Q5 high-fidelity DNA polymerase, NEB; primers TSO_3 and TSO_4, Supplementary Data File 5). The products were purified from a 1% agarose gel (QIAquick Gel Extraction Kit, Qiagen) and ligated by Gibson with 3 h of incubation at 50°C followed by dialysis for 3 h on a membrane filter (MF-Millipore 0.025 μm membrane, Merck) and vacuum concentration. The resulting (NNK1-4) libraries were transformed into 10-beta Electrocompetent E. coli (NEB), by electroporation with 2.0 kV, 200 Ω, 25 μF (BioRad GenePulser machine). Cells were recovered in SOC medium for 30 min and grown overnight in 50 ml of LB ampicillin medium. A small amount of cells was also plated on LB ampicillin plates to assess transformation efficiency. Total transformants were estimated (Supplementary Data File 6), 50 ml of overnight culture were harvested to purify each library with a midi prep (Plasmid MIDI Kit, Qiagen). Libraries NNK1-4 were bottlenecked to ~1 million transformants, while for the replication library we estimated 625,000 transformants.

### Large-scale yeast transformation of random libraries

*Saccharomyces cerevisiae* GT409 [psi-pin-] (MATα ade1–14 his3 leu2-3,112 lys2 trp1 ura3–52) provided by the Chernoff lab was used in all experiments in this study^26^. Yeast cells were transformed with the above plasmid library midipreps. After an overnight pre-growth culture in 25 ml of YPDA medium at 30°C, cells were diluted to OD600 = 0.3 in 175 ml YPDA and incubated at 30°C 200 rpm for ∼4 hr. When cells reached the exponential phase, they were harvested, washed with milliQ, and resuspended in sorbitol mixture (100 mM LiOAc, 10 mM Tris pH 8, 1 mM EDTA, 1M sorbitol). After a 30 min incubation at room temperature (RT), 4 µg of plasmid library and 175 µl of ssDNA (UltraPure, Thermo Scientific) were added to the cells. PEG mixture (100 mM LiOAc, 10 mM Tris pH 8, 1 mM EDTA pH 8, 40% PEG3350) was also added and cells were incubated for 30 min at RT and heat-shocked for 15 min at 42°C in a water bath. Cells were harvested, washed, resuspended in 250 ml recovery medium (YPD, sorbitol 0.5M, 70 mg/L adenine) and incubated for 1.5 hr at 30°C 200 rpm. After recovery, cells were resuspended in 350 ml -URA plasmid selection medium and allowed to grow for 50 hr. Transformation efficiency was calculated for each of the four transformations by plating an aliquot of cells in -URA plates (Supplementary Data File 6). Two days after transformation, the culture was diluted to OD600 = 0.08 in 500 ml -URA medium and grown until exponential phase. At this stage, cells were harvested and stored at -80°C in 25% glycerol. In yeast, libraries NNK1-4 were bottlenecked to 0.5-1 million transformants (Supplementary Data File 6).

### Small-scale yeast transformation of replication library

Yeast cells were transformed with the library containing 400 sequences in three biological replicates. An individual colony was grown overnight in 3 ml YPDA medium at 30 °C and 4 g. Cells were diluted in 60 ml to OD600 = 0.25 and grown for 4–5 h. When cells reached the exponential phase (OD∼0.7–0.8), cells were harvested at 400 × g for 5 min, washed with milliQ, and resuspended in 1 ml YTB (100 mM LiOAc, 10 mM Tris pH 8.0, 1 mM EDTA). They were harvested again and resuspended in 72 µl YTB. 100 ng of plasmid library were added to the cells, together with 8 µl of salmon sperm DNA (UltraPure, Thermo Scientific) previously boiled, 60 µl of dimethyl sulfoxide (Merck) and 500 µl of YTB-PEG (100 mM LiOAc, 10 mM Tris pH 8.0, 1 mM EDTA, 40% PEG 3350). The cells incubated at room temperature for 30 minutes at 4g. Heat-shock was performed at 42 °C for 14 min in a thermo block. Finally, cells were harvested and resuspended in 50 ml plasmid selection medium (-URA, 20% glucose), allowing them to grow for 50 h at 30 °C and 4 g. A small amount of cells was also plated in plasmid selection medium to assess transformation efficiency. We estimated 70,000 transformants per replicate (Supplementary Data File 6). Two days after transformation, the culture was diluted to OD600 = 0.08 in 500 ml -URA medium and grown until exponential phase. At this stage, cells were harvested and stored at -80°C in 25% glycerol.

### Selection experiments

Cells were thawed from −80 °C in 50 ml plasmid selection medium at OD = 0.05 and grown until exponential for 15 h. At this stage, cells were harvested and resuspended in 300 ml protein induction medium (-URA, 2% glucose, 100 μM Cu2SO4) at OD = 0.1. After 24 h the 250 ml input pellets were collected, and cells were plated on -ADE-URA selection medium in 145-cm2 plates (Nunc, Thermo Scientific). Plates were incubated at 30 °C for 7 days. Finally, colonies were scraped off the plates with PBS 1x and harvested by centrifugation to collect the output pellets. Both input and output pellets were stored at −20 °C before DNA extraction. For each random library experiment, one input sample and three technical replicates of the output pellet were processed for sequencing. Selection experiments for the replication library were instead performed in three biological replicates, following the same steps as above. Three input and three output samples were processed for sequencing.

### DNA extraction and sequencing library preparation

Input and output pellets were thawed and resuspended in 1.5 ml extraction buffer (2% Triton-X, 1% SDS, 100 mM NaCl, 10 mM Tris pH 8, 1 mM EDTA pH 8), and underwent two cycles of freezing and thawing in an ethanol-dry ice bath (10 min) and at 62°C (10 min). Samples were then vortexed together with 1.5 ml of phenol:chloroform:isoamyl 25:24:1 and 1.5 g of glass beads (Sigma). The aqueous phase was recovered by centrifugation and mixed again with 1.5 ml phenol:chloroform:isoamyl 25:24:1. DNA precipitation was performed by adding 1:10 V of 3M NaOAc and 2.2 V of 100% cold ethanol to the aqueous phase and incubating the samples at -20°C for 1 hr. After a centrifugation step, pellets were dried overnight at RT. Pellets were resuspended in 900 µl resuspension buffer (10 mM Tris pH 8, 1 mM EDTA pH 8) and treated with 7.5 ml RNase A (Thermo Scientific) for 30 min at 37°C. The DNA was finally purified using 30 µl of silica beads (QIAEX II Gel Extraction Kit, Qiagen), washed and eluted in 22 µl of elution buffer. Plasmid concentrations were measured by quantitative PCR with SYBR green (Merck) and primers annealing to the origin of replication site of the PCUP1-Sup35N plasmid at 58 °C for 40 cycles (TSO_05 and TSO_06, Supplementary Data File 5). The library for high-throughput sequencing was prepared in a two-step PCR (Q5 high-fidelity DNA polymerase, NEB). In PCR1, 160 million of molecules were amplified for 15 cycles at 68°C with frame-shifted primers with homology to Illumina sequencing primers (primers TSO_7 to TSO_20, Supplementary Data File 5). The products were purified with ExoSAP treatment (Affymetrix) and by column purification (MinElute PCR Purification Kit, Qiagen). They were then amplified for 10 cycles in PCR2 with Illumina-indexed primers (primers TSO_21 to TSO_54, Supplementary Data File 5). The library was sequenced by 150 bp paired-end sequencing in an Illumina NextSeq500 sequencer at the CRG Genomics core facility.

### Sequence data preprocessing

We processed each of the 4 NNK experiments separately using DiMSum^58^. Briefly, DiMSum comprises an end-to-end pipeline for processing deep mutational scanning datasets from raw reads to measured sequences and their associated assay scores (plus errors). DiMSum was run with the following parameters: cutadaptMinLength=“60”; cutadaptErrorRate=“0.2”; vsearchMinQual=“30”; vsearchMaxee=“0.5”; startStage=“0”; fitnessMinInputCountAny=“0”; maxSubstitutions=“20”; mixedSubstitutions=“TRUE”; experimentDesignPairDuplicates=“TRUE”. We then removed sequences with fewer than 100 reads in the input sequencing experiment. Next, we centered the fitness estimates (nucleation scores) of each dataset individually by adding or subtracting the corresponding mode fitness of the non-nucleating sequences. After centering each sequence, we next labeled sequences as “nucleators” (or “non-nucleators”) by transforming their fitness estimate to a Z-score composed of the fitness estimate scaled by the DiMSum error, and performing a one-sided hypothesis test to check whether standardized score was significantly larger than 0. We treated sequences whose p-values after FDR adjustment were <= 0.05 as “nucleators”, and remaining sequences as “non-nucleators.” A proportion of sequences produced no reads after the selection experiments, thus leading to NA scores from DiMSum. We labeled these sequences as “non-nucleators.” If a sequence contained a stop codon, we used only the component of the sequence preceding the stop for model training. For cases in which this resulted in duplicate sequences (e.g. FN*VILRDEGHGSYGFDNNN and FN*FVVMHTCIMVVFCLGDI are both mapped to “FN”), we summarized the truncated sequence by taking its mean nucleation score or mode nucleation status across observations. If a given truncated sequence had an equal number of nucleator and non-nucleator status observations, we discarded this truncated sequence. As a result we classified > 35,000 sequences for libraries NNK1-3 (35,456; 37,578; 38,893 respectively) and 7,040 for NNK4.

### The architecture of CANYA

CANYA is a biologically motivated hybrid-neural network designed to discover motifs and their interactions. More concretely, the architecture of CANYA is inspired by recent work that suggests stacked convolution and attention layers serve as a reasonable inductive bias for motif and motif-interaction discovery. The hyperparameters of CANYA were influenced by summary statistics of interacting secondary structure elements in amyloids within the PDB (Supplementary Fig. 4). Summarily, we chose the simplest architecture of our model such that it is expressive, interpretable, and importantly, principled in biological knowledge.

CANYA takes as input an amino acid sequence of length limit up to 145 residues, and outputs a score related to the sequence’s propensity to form amyloids. Prior to passing the sequence to the input layer, we first one-hot encode it, allowing only the 20 canonical amino acids. As we use filters of length 3 (See below for justification; Supplementary Fig. 4), we pad the sequence with two 0s both up- and downstream the sequence. Finally, if this padded sequence is not of length 149, we add a mask with values of -1 downstream the sequence until it reaches length 149. The input length restrictions of CANYA arise from the fact that a given sequence in the assay is fused to a Sup35N construct of length 125, is (up to) length 20, and is padded with two 0s on each side. Explicitly, the training data of CANYA looks as follows:

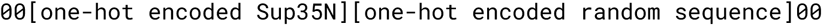

when there is no masking or stop codons, and as follows if so:

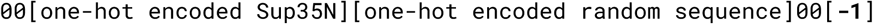

where the number of -1 values is the required quantity such that the sequence is length 149.

The input layer of CANYA correspondingly accepts a matrix of size 149×20 representing a one-hot encoded, padded, and potentially masked peptide sequence. The output layer is a single unit with sigmoid activation. The hidden layers of CANYA are:

1. Convolution (100 filters, size 3, stride 1, exponential activation)
2. Self-attention (1 attention head, key-length 6)
3. Fully-connected layer (64 units, ReLU activation)

We selected an exponential activation function for the convolutional layer as this type of activation is generally more robust for motif discovery^59^. We chose filters of length 3 as this was the mode length of beta-sheets in amyloid sequences with resolved structures in Uniprot (Supplementary Fig. 4). We utilize dropout with probability 0.1 after the convolution and attention layers, and 0.4 after the fully-connected layer. We use an elastic net regularization (with value 0.01) when learning the weights between the attention and fully-connected layers. Finally, to encourage the model to learn positional information, we do not perform pooling after the convolution layer, and we include positional encodings prior to taking the softmax in the attention layer. We trained CANYA for 100 epochs using the adam optimizer with default values and the binary Kullbeck-Leibler divergence as a loss function. We limited the learning rate of the model during training by monitoring the validation area under precision-recall curve, decaying at a factor of 0.2 with patience 4, and performed early stopping by monitoring the validation area under precision-recall curve with patience 10. For sequences with length greater than 145, we collect the CANYA score at every overlapping length-145 window of the sequence, then use its minimum CANYA score as its final score (under the logic that nucleation-forming propensity is limited by a sequence’s most nucleation-disrupting region).

### Compilation of external datasets

We first collected 6-mers from the WALTZ-DB dataset^28^. Here, we assigned all sequences whose “Classification” field was “amyloid” as a 1, and all other sequences as 0. We next collected the collection of aggregating peptides from the CPAD repository^43^. We used sequences from the “Peptide” field, filtering for sequences of at least length 10 and for sequences that did not contain a space in their sequences. We assigned sequences with “Classification” field “Amyloid” a 1, and all other sequences 0. We then collected the Amy dataset from the AMYPred-FRL server^23^. The Amy database contains literature-mined amyloid precursor proteins with validated amyloidogenic sequence regions—portions of an amyloid-forming protein that when isolated from the remaining sequence have been confirmed to form amyloids in external experiments. The median length of sequences in Amy (539) is substantially longer than those of the previous datasets (next longest median length=19 in NNK4, or 16 in CPAD), so this prediction task evaluates whether methods can account for context-specific and distal effects when generating their propensity scores. Here, we assigned all sequences in the negative sets a label of 0, and all sequences in the positive sets a label of 1. Many of the sequences had lengths greater than 145, we therefore applied a sliding window approach to these sequences in which we score every overlapping length-145 region of a sequence, and assigned a final score to the entire sequence as the minimum of the length-145 regional scores. The final external dataset we used was from the AmyPro database^34^. Though AmyPro contains overlapping sequences with the Amy dataset, we treated this task differently than the previous tasks. Namely, all sequences in the AmyPro dataset were amyloids, and so we sought to evaluate methods’ abilities to distinguish the amyloidogenic region from the non-amyloidogenic regions of the sequences. First, we collected all sequences from the “regions” field in the dataset. Next we removed each of these “region” sequences from the main peptide sequence and concatenated the remaining two portions of the main sequence together, comprising a set of positive sequences (labeled 1) from the “region” field and negative sequences (labeled 0) from the remaining peptide sequences. Finally, we limited the length of all sequences to 100 by breaking sequences longer than length 100 into non-overlapping subsequences of at most length 100. While this task evaluates un-natural sequences, it evaluates the ability of each method to distinguish amyloid cores from non-amyloid cores while also making the problem more amenable to previous approaches, which generally underperformed on long sequences. We list descriptive summary statistics (e.g. length, sample sizes, hydrophobicity) in Supplementary Table 5.

### Aggregation predictors

Aggregation predictors or physicochemical scales (Tango^12^, Amypred^23^, Camsol^13^, PLAAC^14^, Aggrescan^15^) were used to calculate a score for each sequence. When appropriate, individual residue-level scores were summed to obtain a single score per sequence. CamSol, Amypred and Aggrescan were run with the default parameters. PLAAC was run using a core of length 6 and weightings from input sequences. Tango was run with pH 7.2, no protection of termini, ionic strength = 0.1 and T = 298K (25°C). Some of the predictors present sequence length limitations: Amypred runs only for sequences longer than 10 amino acids, CamSol for sequences longer than 6 amino acids, and Aggrescan cannot be run for sequences longer than 2004 amino acids. We note that several of these methods (including TANGO, CamSol, and Aggrescan) were directly trained on sequences within CPAD, and other datasets presented in the manuscript, violating the ability to evaluate their out-of-sample predictive performance on these datasets. This complication is exacerbated by several methods (e.g., TANGO, CamSol) also being ensemble methods (or extensions) that leverage several algorithms for prediction—it is not trivial to account for, or remove, these previously seen sequences, as any sequence that was used for training the main algorithm or their antecedent ensemble methods is not out-of-sample.

### Selecting a model for interpretability analyses

We trained CANYA with random weight initialization 100 times and recorded for each fitted model the area under the curve (AUC) of the test data, area under the precision-recall curve (AUPROC) of the test data, and interpretability score adapted from a recently developed approach for interpretability analyses of genomic neural networks^31^. Briefly, Majdandzic et al. propose an approach to quantify the consistency of the attribution maps of a trained model by comparing the entire set of kmers in the training sequences to the set of kmers in (adjusted^44^) attributed positions in the training sequences. These two distributions of kmers—in the case of CANYA, 3-mers—are compared using the Kullbeck-Leibler (KL) divergence, where a higher KL divergence suggests greater amenability to downstream interpretability analyses. To calculate an interpretability score for each trained instance of CANYA, we used this same approach, but rather than using kmers of nucleotides, we used kmers from the input amino acids. As we saw that the test AUPROC was more consistent across experiments, we used a models’ mean AUPROC across experiments and interpretability score as model selection criteria. More rigorously, we selected the model with the highest interpretability score, conditional on the fact that its mean AUPROC across datasets was greater than the median of these mean scores across model training instances.

### Visualization of filters (motifs)

Notably, the use of random sequences in amino acid space poses difficulties for observing a typical, lexicographic motif, and consequently, observing convergence toward a lexicographic motif in first-layer convolutional filters. We elaborate as follows: using a filter length of 3, there is a 1 in 8,000 (20^3^) chance of observing a given kmer. Ideally, for the model to learn a stable feature, this kmer must not only exist in a sizable proportion of sequences, but its effect must also not be masked out by surrounding contextual information. Even if we were to ignore contextual information, this motif would need to occur independently multiple times, an event whose probability quickly converges to 0. Consequently, we are much stricter than previous approaches when generating a position weight matrix (PWM) for a given filter. For interpretability’s sake, we limit the kmers comprising a PWM for a filter to the minimum of either the 10 most-activating kmers of a filter, or the collection of kmers whose activation is at least 75% of the maximum-activating kmer. Summarily, a filter is both visualized and represented numerically by its PWM composed of at most the top 10 strongest activating kmers.

### Motif clustering

Following the above logic, CANYA must learn physicochemical properties of amino acids and understand how these properties interact amongst each other when constructing its features at the convolution layer. Moreover, these physicochemical 3-mers, or motifs, may often capture redundant physicochemical information, but independent sequences—for example, two different motifs capturing hydrophobicity may separately comprise sequences of “IVF” or “ALM.” To further improve interpretability and reduce the dimensionality of downstream experiments leveraging the learned motifs of CANYA, we performed clustering on the PWM matrices. More concretely, we calculated BLOSUM scores for each filter by taking the dot product between its PWM and BLOSUM score matrix^39^. We next performed affinity propagation on these calculated motif BLOSUM scores to cluster the motifs. Affinity propagation discovered 10 clusters of motifs. However, after performing Global Importance Analysis (GIA) experiments^35^, we found 7 discrepancies when evaluating whether a given motif had the same effect size (importance score) direction compared to the effect size of the motif with the greatest absolute effect within the cluster. As our goal was to interpret model decisions and physicochemical clusters, we removed these 7 filters from their corresponding clusters so that each cluster contained only filters with the same effect size direction. We show the original and changed cluster assignments in Supplementary Figure 6.

### Global Importance Analysis (GIA) Experiments

To learn the effect of motif presence on CANYA’s decision-making, we turned to Global Importance Analysis (GIA) *in-silico* experiments^35^. Briefly, GIA is a post-hoc interpretability method applied to genomic neural networks that enables users to learn importance scores (i.e. effect sizes) of a given sequence feature on a model’s output score. The importance score is derived from taking the average difference in model score between a set of background sequences, and this same set of background sequences but with a functional element, such as a motif, placed in the background sequence (sequence length is maintained, i.e. a window of the sequence is replaced by the functional element). For all experiments, we limited our analyses to 25,000 randomly selected, full-length (length-20 and and absent of stop codons) training sequences that were confidently predicted by CANYA. We defined “confidently predicted” as nucleators with CANYA score above 0.3 and non-nucleators with CANYA score below 0.2 (see Supplementary Fig. 9 for prediction score distributions). Finally, we emphasize that owing to the random nature of our experiment, the training sequences serve as a valid set of background sequences for GIA as they span an extremely wide range of contexts.

In the first set of GIA experiments, we sought to characterize the importance score of each filter individually. To do so, we first randomly selected 25,000 sequences from the training set, comprising sequences from across all three experiments. Next, for a given filter, we collected the activation energy of each kmer used to represent the PWM, and used the ratio of the activation energy of each kmer to the activation energy of the kmer with the maximum activation energy in this PWM to generate kmer sampling probabilities. For each sequence, we randomly sampled one kmer using this normalized ratio as the kmer’s sampling probability, and embedded this kmer into the sequence. Afterward, we calculated for all 25,000 background sequences and all 25,000 modified sequences the CANYA nucleation score prior to applying the softmax function. We calculated each filter’s importance score as the mean paired difference in scores between the 25,000 background and modified sequences.

After clustering the learned motifs, we next wished to validate whether the clusters could be utilized to simplify further interpretability analyses by reducing the scale of *in-silico* experiments performed. To do so, we conducted a GIA experiment within each cluster to determine a cluster-level importance score. The experiment follows the same logic as the original, filter-level GIA experiment, only that we first randomly selected a filter within a cluster prior to sampling a kmer from its PWM. The filters were randomly selected according to the ratio of their absolute GIA importance score to the maximum absolute GIA importance score across filters of the corresponding cluster. Indeed, cluster-level scores recapitulated the scores of the motifs from which they were composed (Supplementary Table 5). We therefore performed all following GIA analysis at the cluster level, using this filter-first, kmer-second sampling scheme.

We next performed an experiment to evaluate the additivity of motif-clusters on nucleation propensity. Here, we collected 25,000 background sequences from the training dataset, then embedded into these background sequences 1 to 4 kmers in non-overlapping positions where each of the 4 kmers was sampled using the filter-first, kmer-second sampling scheme. Each sequential kmer addition (from kmers 2-4) was embedded in the sequence such that the sequence with antecedent kmer multiplicity maintained the kmer(s) at its (their) original embedded position(s). We calculated the cluster importance score for a given multiplicity by taking the mean difference in prediction score between the sequences with the injected kmer(s) and their corresponding background sequences—in other words, each importance score is generated by taking the mean difference between 25,000 background sequences and 25,000 modified background sequences with either 1, 2, 3, or 4 embedded kmers.

To evaluate whether CANYA learned position-specific importance of motifs, we performed an additional GIA experiment in which we systematically embedded a motif-cluster at each position of a random sequence. In these experiments, we performed a single GIA experiment with 25,000 background sequences and 25,000 modified sequences for each position from positions 1-18, so that the entire 3-mer could be contained within the sequence.

In a final GIA experiment, we characterized interaction effects between motif-clusters. For a given motif-cluster pair, we sampled a kmer (as mentioned above) from each cluster as well as a corresponding position randomly from positions 1-18 in which to embed each kmer. We evaluated the CANYA score for the background sequence, the background sequence with the kmer from the first cluster at the first sampled position, the background sequence with the kmer from the second cluster at the second sampled position, and the background sequence with both kmers at both positions. We called the interaction importance as the result of subtracting the sum of CANYA predictions of the sequences with each marginal kmer embedding from the sum of the CANYA predictions of the background sequence and sequence with both motifs. The final importance was calculated as the mean interaction importance across 25,000 sequences.

### Secondary structure enrichment scoring of motifs

To examine whether certain motifs were characteristically similar to sequences found in specific secondary elements of amyloids, we examined activation energies of filters across secondary structure elements in a set of amyloids with resolved structures in the PDB. Concretely, we collected 114 entries from the STAMP dataset^40^, then downloaded their structural information from the PDB (see Supplementary Table 7 for entries and corresponding proteins). Next, we passed all sequences through CANYA, and extracted their filter activation energies (i.e., output from the convolution layer). At each position, we summarized a cluster’s activation energies as the maximum activation energy across filters within a cluster, generating a vector of maximum activation energies for each cluster. Next, we encoded each secondary structure (coil, beta strand, or disorder) as a binary vector where 1 indicated positions in the corresponding secondary structure, and 0 indicated otherwise. We collected this set of secondary structure vectors and activation energy vectors for all sequences, then concatenated them across sequences. Finally, we generated secondary structure enrichment scores by calculating the AUC between a given secondary structure element and cluster activation energy across all sequences.

## Supplementary Data

**Supplementary Data File 1** All sequences recorded spanning each experiment with reported fitnesses, error, and nucleation status

**Supplementary Data File 2** Sequences used to train and test CANYA

**Supplementary Data File 3** Sequences used in replication experiments with their original measured fitness and fitness from the replication experiment

**Supplementary Data File 4** Validation sequences and their corresponding nucleotide sequences

**Supplementary Data File 5** Oligo pool and primer sequences for the NNK experiments

**Supplementary Data File 6** Transformants measured across each experiment

---

**Supplementary Figure 1.**
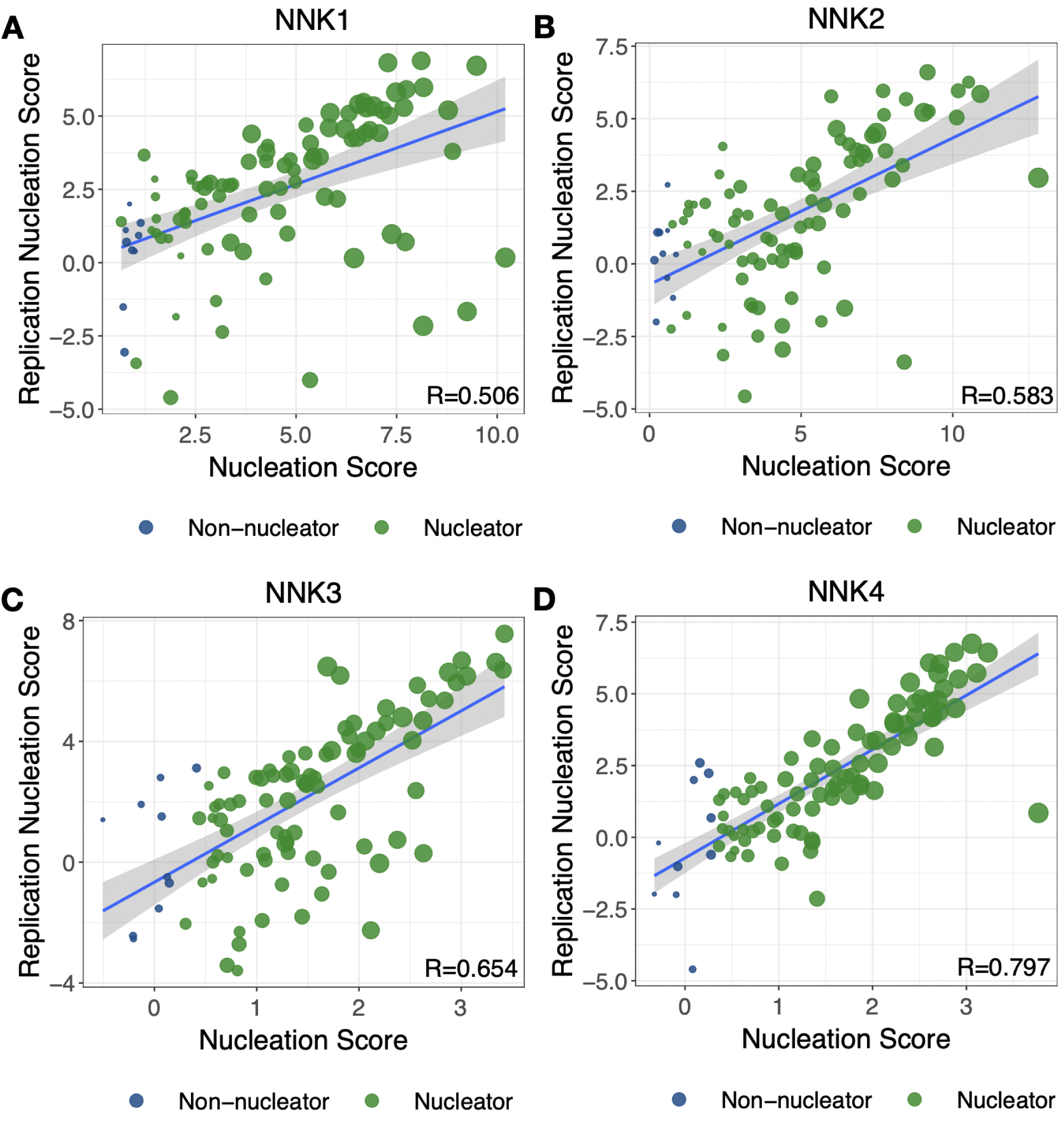
Nucleation scores are replicable across experiments. Nucleation scores from the corresponding original experiment are plotted on the x-axis, and the replication nucleation score is plotted on the y-axis. Sizes are proportional to the inverse error measurement from the original experiment as reported by DiMSum. The x-axis scores were all calculated independently within their respective experiment—(A) NNK1, (B) NNK2, (C) NNK3, (D) NNK4, the validation set—and altogether in the replication set (n=100), as 100 sequences were taken from each experiment.

**Supplementary Figure 2.**
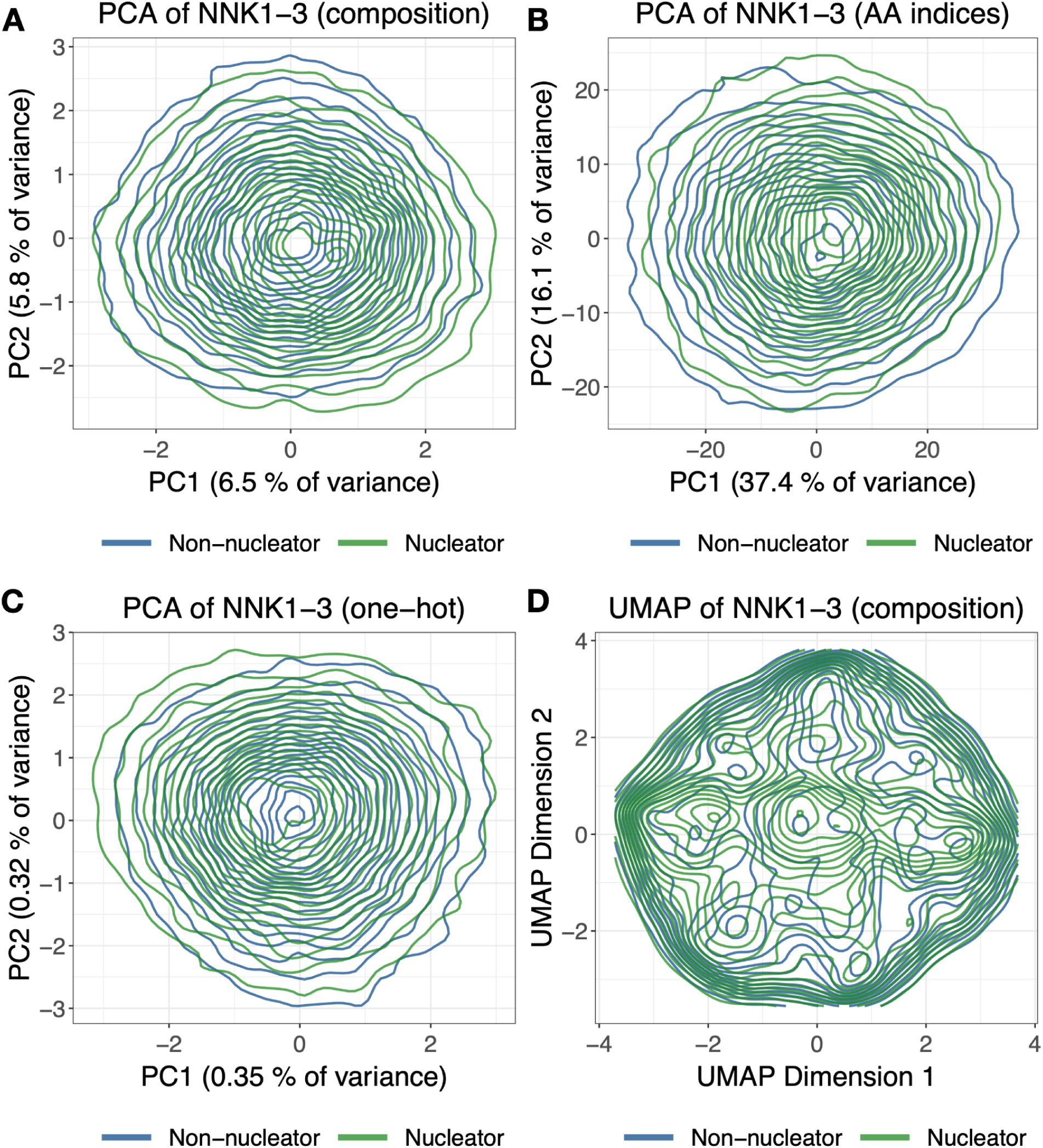
Dimensionality techniques fail to distinguish nucleation status. (A-C) Contour plots of Principal Component Analysis (PCA) scores on PCs 1 and 2 when sequences are represented by (A) overall amino acid composition, (B) 533 amino acid indices calculated from python package protlearn (C) one-hot (position-maintained) amino acid composition. (D) UMAP projection when using amino acid composition as input.

**Supplementary Figure 3.**
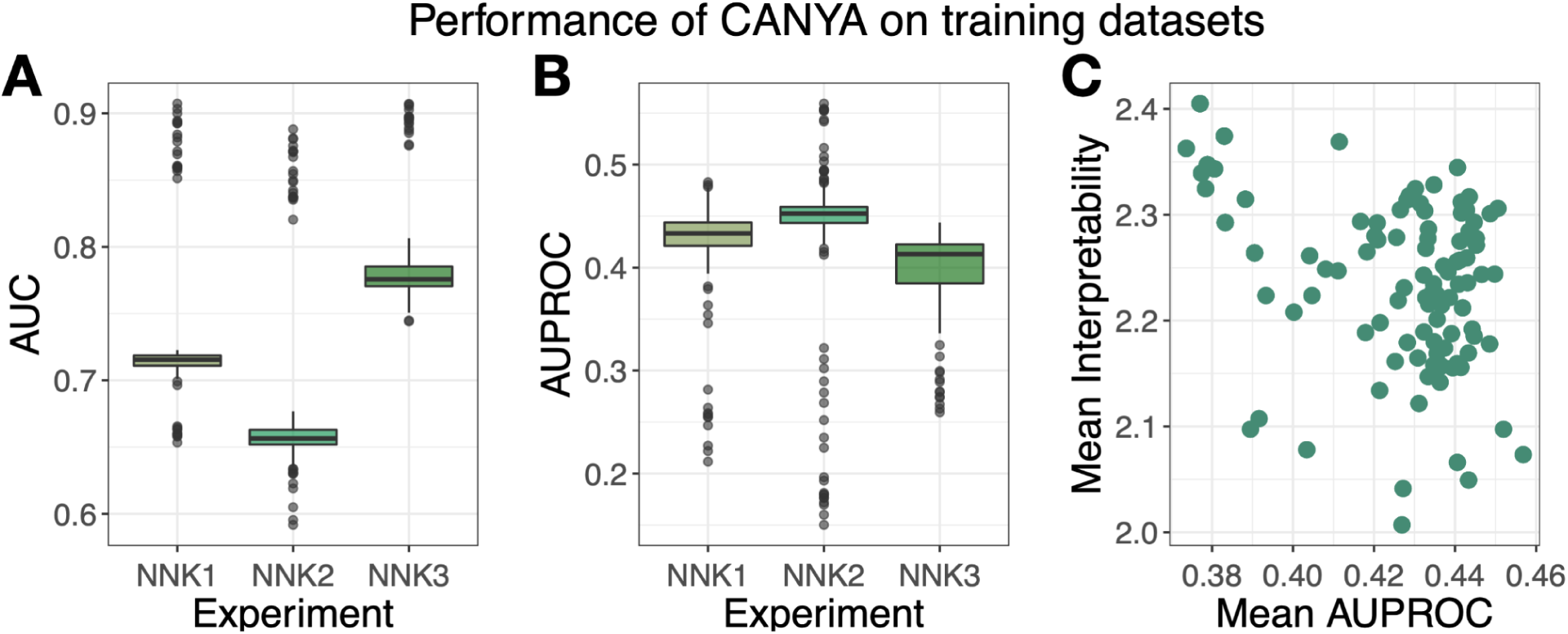
CANYA performance on the held-out test portion of the training datasets. Evaluation metrics across the all 100 model fits of CANYA. (B) The area under receiver operating characteristic curve (AUC) for held-out testing sequences. (C) The area under precision recall curve (AUPROC) for held-out testing sequences. (D) The interpretability score (KL divergence; Methods) calculated on all held-out test sequences plotted against the mean AUPROC across experiments.

**Supplementary Figure 4.**
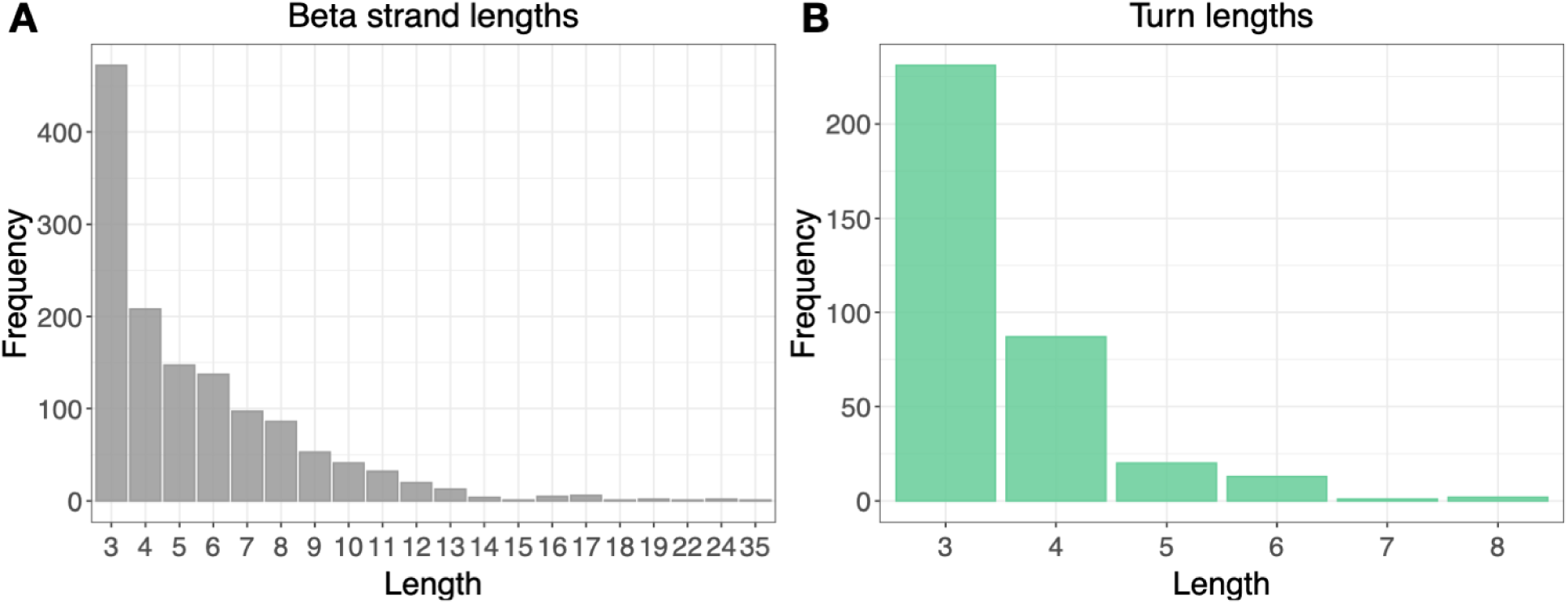
Secondary structure lengths of resolved amyloids. We downloaded data from the WALTZ-DB data portal, filtered for amyloids, then manually parsed the Uniprot entries of each sequence to obtain the distribution of (A) beta strands and (B) turns across 80 sequences.

**Supplementary Figure 5.**
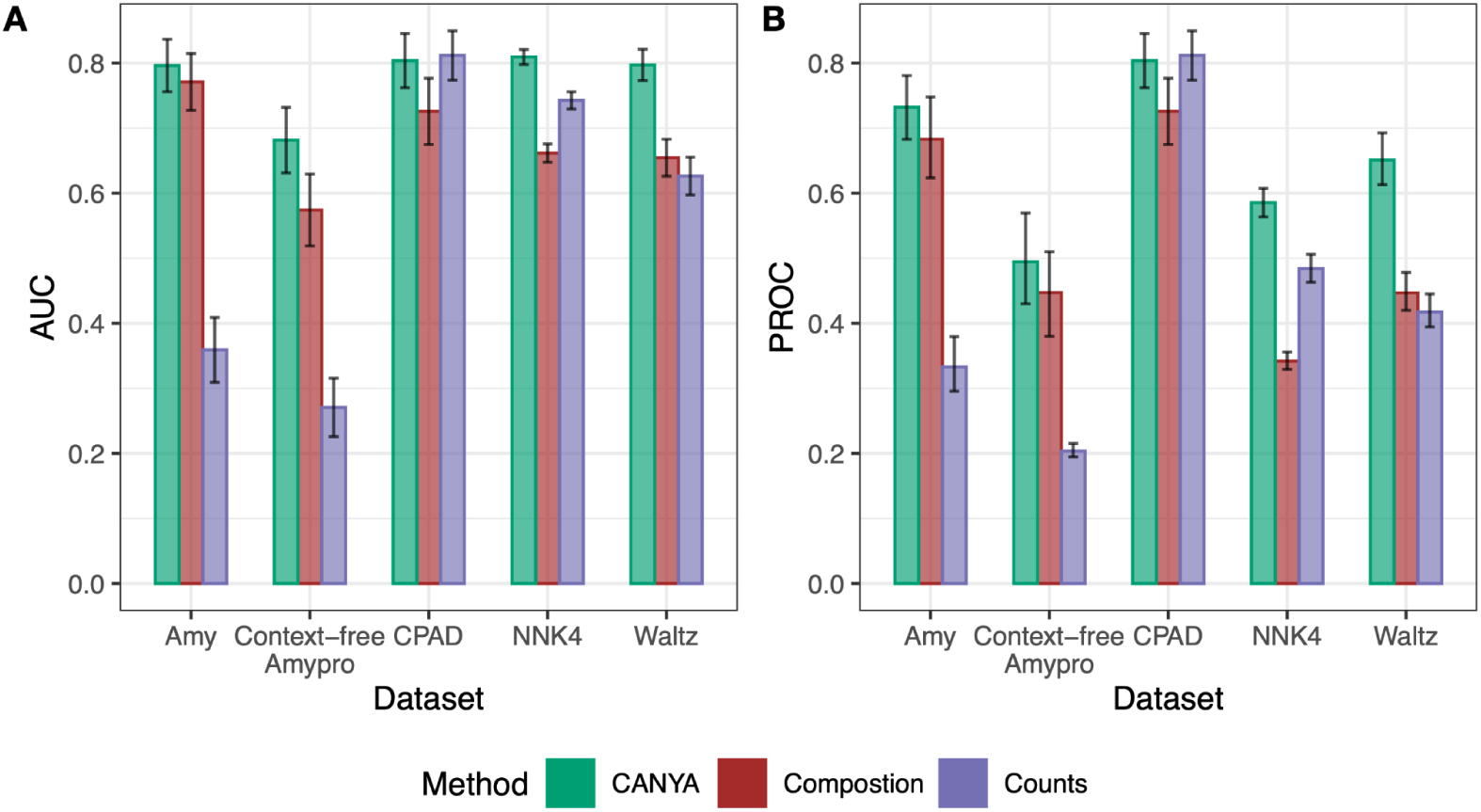
The performance of CANYA compared to simpler models. Area under the receiver-operating characteristic and precision-recall curves (AUC, AUPROC respectively) of each method on a corresponding testing set (Methods, Supplementary Table 6). “Composition” corresponds to training a simple logistic regression model using amino acid composition (between 0.0 and 1.0, proportions) over the training NNK dataset, and “Counts” corresponds to training the same model with raw, unnormalized amino acid counts (between 0 and 20, integer values).

**Supplementary Figure 6.**
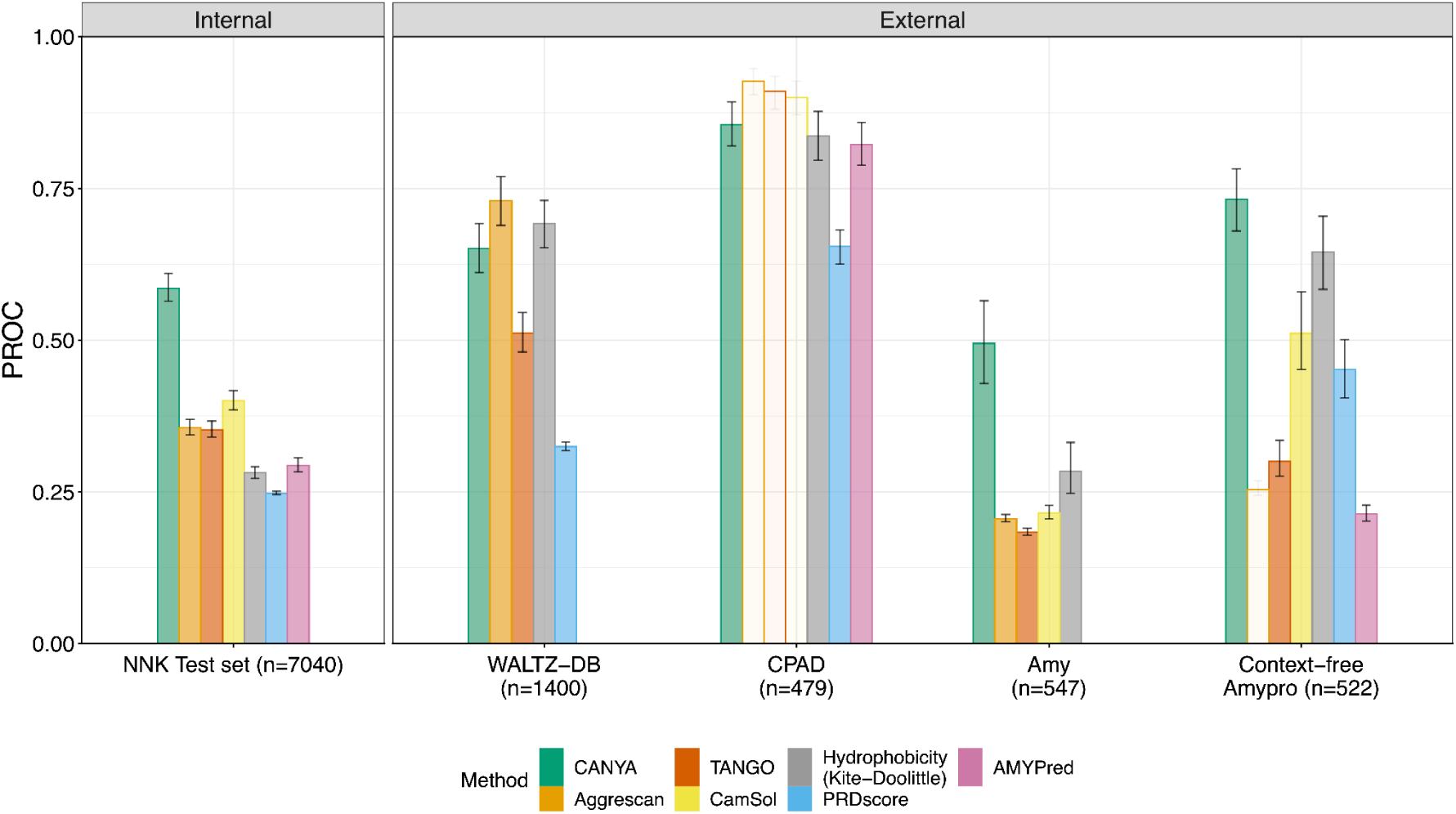
The performance of CANYA compared to previous approaches on testing datasets. Area under the precision-recall curve (AUPROC) of each method on a corresponding testing set. Low-opacity bars represent cases in which the method used data from the testing dataset to do its training, and thus are not valid out-of-sample evaluations. See text for additional descriptions of datasets (Methods, Supplementary Table 6).

**Supplementary Figure 7.**
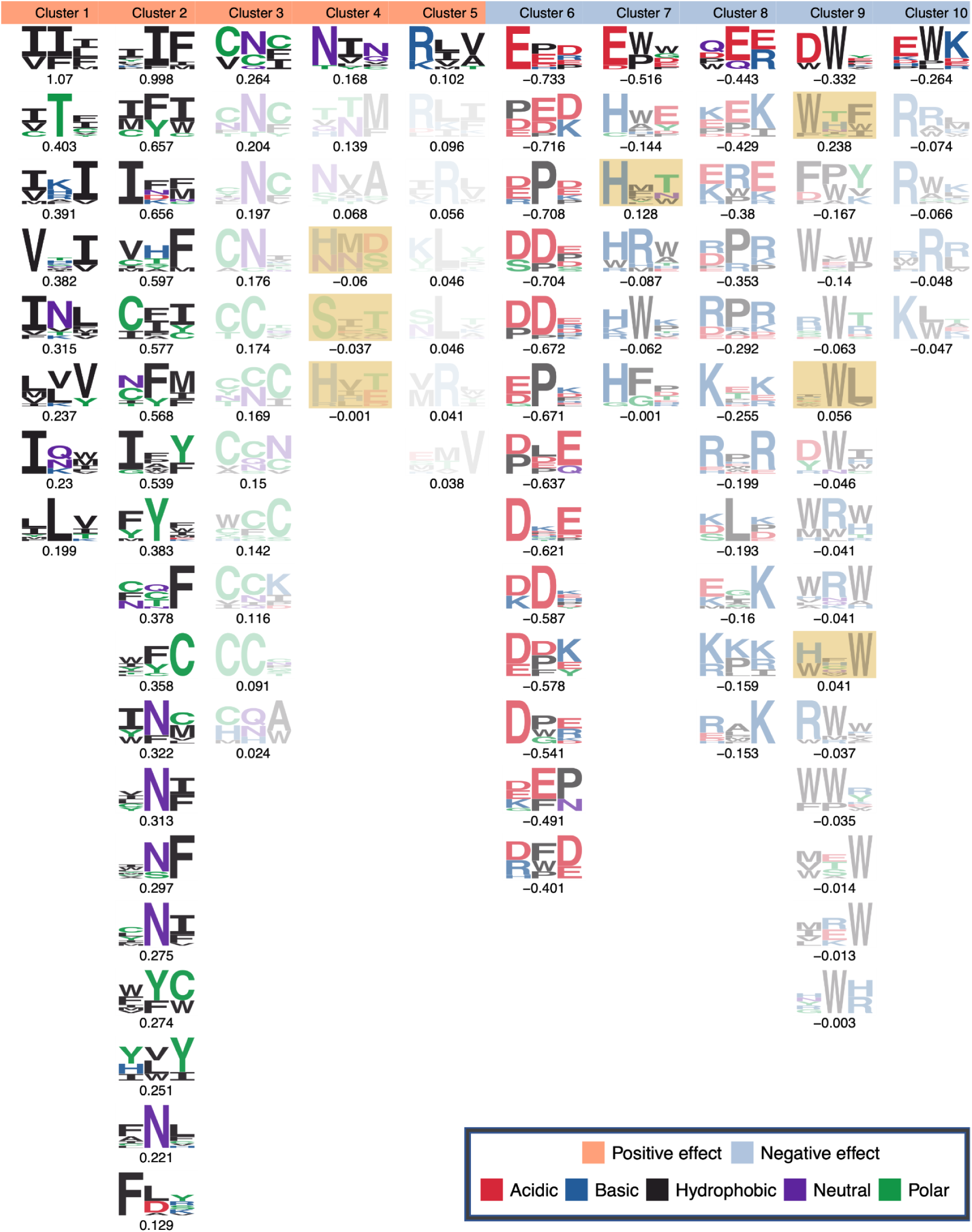
Physicochemical motifs discovered by CANYA prior to performing quality control. We removed filters from clusters if their GIA effect direction was opposite the sign of the effect of the strongest filter. We highlight which motifs were excluded from downstream xAI analysis in yellow.

**Supplementary Figure 8.**
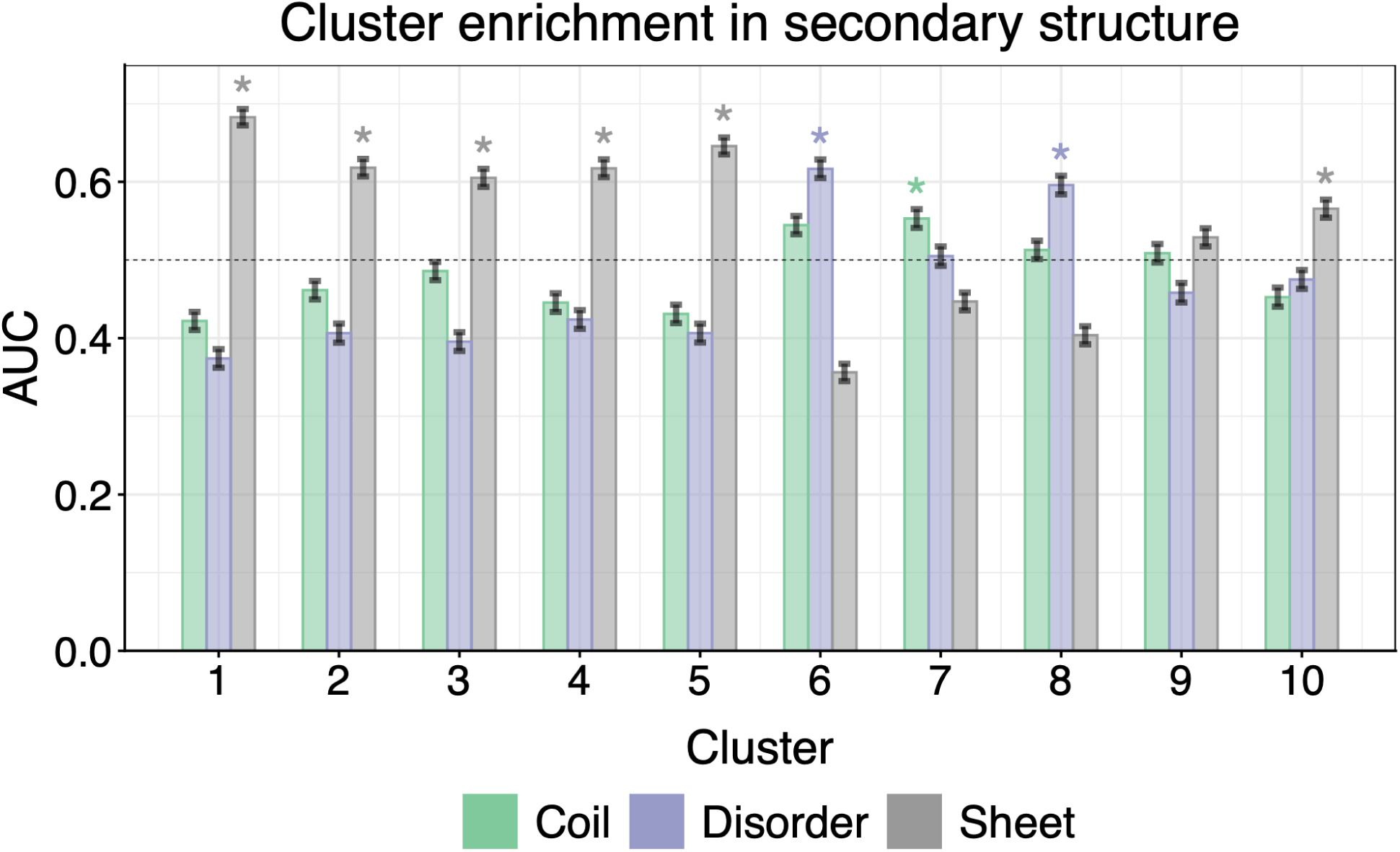
Secondary structure enrichment of motifs discovered by CANYA. We collected sequences from the StAmP dataset then collected their convolution layer activation energies from CANYA. Across all sequences, we examined whether a specific cluster had higher activation (pattern matching) within a specific secondary structure by calculating the AUC between the activation energy on a specific secondary structure (Methods). Asterisks represent structures for which the enrichment was significantly higher than both 0.50 and the second most-enriched structure.

**Supplementary Figure 9.**
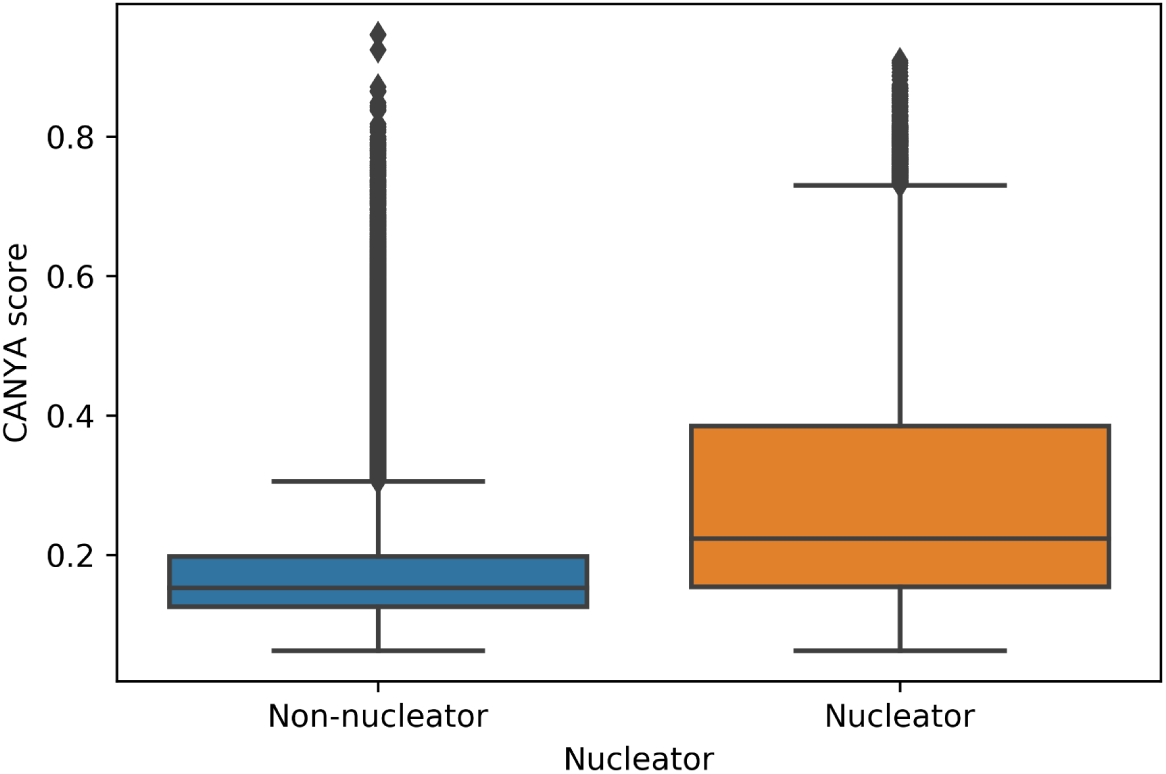
Distribution of CANYA scores across training sequences. The distribution of output scores for non-nucleators (n=79,910) and nucleators (n=20,826).

## References

1. Chiti, F. & Dobson, C. M. Protein Misfolding, Amyloid Formation, and Human Disease: A Summary of Progress Over the Last Decade. Annu. Rev. Biochem. 86, 27–68 (2017).

2. Fowler, D. M., Koulov, A. V., Balch, W. E. & Kelly, J. W. Functional amyloid--from bacteria to humans. Trends Biochem. Sci. 32, 217–224 (2007).

3. Shire, S. J. Formulation and manufacturability of biologics. Curr. Opin. Biotechnol. 20, 708–714 (2009).

4. Ke, P. C. et al. Half a century of amyloids: past, present and future. Chem. Soc. Rev. 49, 5473–5509 (2020).

5. Dobson, C. M., Knowles, T. P. J. & Vendruscolo, M. The Amyloid Phenomenon and Its Significance in Biology and Medicine. Cold Spring Harb. Perspect. Biol. 12, (2020).

6. Scheres, S. H. W., Ryskeldi-Falcon, B. & Goedert, M. Molecular pathology of neurodegenerative diseases by cryo-EM of amyloids. Nature 621, 701–710 (2023).

7. Sabaté, R. & Ventura, S. Cross-β-sheet supersecondary structure in amyloid folds: techniques for detection and characterization. Methods Mol. Biol. 932, 237–257 (2013).

8. Eisenberg, D. S. & Sawaya, M. R. Structural Studies of Amyloid Proteins at the Molecular Level. Annu. Rev. Biochem. 86, 69–95 (2017).

9. Shi, Y. et al. Structure-based classification of tauopathies. Nature 598, 359–363 (2021).

10. Yang, Y. et al. Cryo-EM structures of amyloid-β 42 filaments from human brains. Science 375, 167–172 (2022).

11. Schweighauser, M. et al. Structures of α-synuclein filaments from multiple system atrophy. Nature 585, 464–469 (2020).

12. Fernandez-Escamilla, A.-M., Rousseau, F., Schymkowitz, J. & Serrano, L. Prediction of sequence-dependent and mutational effects on the aggregation of peptides and proteins. Nat. Biotechnol. 22, 1302–1306 (2004).

13. Sormanni, P., Aprile, F. A. & Vendruscolo, M. The CamSol method of rational design of protein mutants with enhanced solubility. J. Mol. Biol. 427, 478–490 (2015).

14. Lancaster, A. K., Nutter-Upham, A., Lindquist, S. & King, O. D. PLAAC: a web and command-line application to identify proteins with prion-like amino acid composition. Bioinformatics 30, 2501–2502 (2014).

15. Conchillo-Solé, O. et al. AGGRESCAN: a server for the prediction and evaluation of ‘hot spots’ of aggregation in polypeptides. BMC Bioinformatics 8, 65 (2007).

16. Wickner, R. B. et al. Yeast Prions Compared to Functional Prions and Amyloids. J. Mol. Biol. 430, 3707–3719 (2018).

17. Wickner, R. B. Yeast and Fungal Prions. Cold Spring Harb. Perspect. Biol. 8, (2016).

18. Wilkinson, M. et al. Structural evolution of fibril polymorphs during amyloid assembly. Cell 186, 5798–5811.e26 (2023).

19. Lövestam, S. et al. Disease-specific tau filaments assemble via polymorphic intermediates. Nature 625, 119–125 (2024).

20. Knowles, T. P. J., Vendruscolo, M. & Dobson, C. M. The amyloid state and its association with protein misfolding diseases. Nat. Rev. Mol. Cell Biol. 15, 384–396 (2014).

21. Baldwin, A. J. et al. Metastability of native proteins and the phenomenon of amyloid formation. J. Am. Chem. Soc. 133, 14160–14163 (2011).

22. Navarro, S. & Ventura, S. Computational methods to predict protein aggregation. Curr. Opin. Struct. Biol. 73, 102343 (2022).

23. Charoenkwan, P. et al. AMYPred-FRL is a novel approach for accurate prediction of amyloid proteins by using feature representation learning. Sci. Rep. 12, 7697 (2022).

24. Seuma, M., Faure, A. J., Badia, M., Lehner, B. & Bolognesi, B. The genetic landscape for amyloid beta fibril nucleation accurately discriminates familial Alzheimer’s disease mutations. Elife 10, (2021).

25. Seuma, M., Lehner, B. & Bolognesi, B. An atlas of amyloid aggregation: the impact of substitutions, insertions, deletions and truncations on amyloid beta fibril nucleation. Nat. Commun. 13, 7084 (2022).

26. Chandramowlishwaran, P. et al. Mammalian amyloidogenic proteins promote prion nucleation in yeast. J. Biol. Chem. 293, 3436–3450 (2018).

27. Louros, N., Orlando, G., De Vleeschouwer, M., Rousseau, F. & Schymkowitz, J. Structure-based machine-guided mapping of amyloid sequence space reveals uncharted sequence clusters with higher solubilities. Nat. Commun. 11, 3314 (2020).

28. Louros, N. et al. WALTZ-DB 2.0: an updated database containing structural information of experimentally determined amyloid-forming peptides. Nucleic Acids Res. 48, D389–D393 (2020).

29. Ullah, F. & Ben-Hur, A. A self-attention model for inferring cooperativity between regulatory features. Nucleic Acids Res. 49, e77 (2021).

30. Ghotra, R. S., Lee, N. K. & Koo, P. K. Uncovering motif interactions from convolutional-attention networks for genomics. NeurIPS 2021 AI for Science Workshop (2021).

31. Majdandzic, A. et al. Selecting deep neural networks that yield consistent attribution-based interpretations for genomics. Proc Mach Learn Res 200, 131–149 (2022).

32. Kyte, J. & Doolittle, R. F. A simple method for displaying the hydropathic character of a protein. J. Mol. Biol. 157, 105–132 (1982).

33. Niu, M., Li, Y., Wang, C. & Han, K. RFAmyloid: A Web Server for Predicting Amyloid Proteins. Int. J. Mol. Sci. 19, (2018).

34. AmyPro database.

35. Koo, P. K., Majdandzic, A., Ploenzke, M., Anand, P. & Paul, S. B. Global importance analysis: An interpretability method to quantify importance of genomic features in deep neural networks. PLoS Comput. Biol. 17, e1008925 (2021).

36. Sawaya, M. R., Hughes, M. P., Rodriguez, J. A., Riek, R. & Eisenberg, D. S. The expanding amyloid family: Structure, stability, function, and pathogenesis. Cell 184, 4857–4873 (2021).

37. Murray, K. A. et al. Identifying amyloid-related diseases by mapping mutations in low-complexity protein domains to pathologies. Nat. Struct. Mol. Biol. 29, 529–536 (2022).

38. Kanchi, P. K. & Dasmahapatra, A. K. Polyproline chains destabilize the Alzheimer’s amyloid-β protofibrils: A molecular dynamics simulation study. J. Mol. Graph. Model. 93, 107456 (2019).

39. Henikoff, S. & Henikoff, J. G. Amino acid substitution matrices from protein blocks. Proc. Natl. Acad. Sci. U. S. A. 89, 10915–10919 (1992).

40. Louros, N., van der Kant, R., Schymkowitz, J. & Rousseau, F. StAmP-DB: a platform for structures of polymorphic amyloid fibril cores. Bioinformatics 38, 2636–2638 (2022).

41. Izawa, Y. et al. Role of C-terminal negative charges and tyrosine residues in fibril formation of α-synuclein. Brain Behav. 2, 595–605 (2012).

42. Tompa, P. Structural disorder in amyloid fibrils: its implication in dynamic interactions of proteins. FEBS J. 276, 5406–5415 (2009).

43. Thangakani, A. M. et al. CPAD, Curated Protein Aggregation Database: A Repository of Manually Curated Experimental Data on Protein and Peptide Aggregation. PLoS One 11, e0152949 (2016).

44. Majdandzic, A., Rajesh, C. & Koo, P. K. Correcting gradient-based interpretations of deep neural networks for genomics. Genome Biol. 24, 109 (2023).

45. Li, F.-Z., Amini, A. P., Yue, Y., Yang, K. K. & Lu, A. X. Feature Reuse and Scaling: Understanding Transfer Learning with Protein Language Models. bioRxiv 2024.02.05.578959 (2024) doi:10.1101/2024.02.05.578959.

46. Flamholz, Z. N., Biller, S. J. & Kelly, L. Large language models improve annotation of prokaryotic viral proteins. Nat Microbiol 9, 537–549 (2024).

47. Thumuluri, V., Almagro Armenteros, J. J., Johansen, A. R., Nielsen, H. & Winther, O. DeepLoc 2.0: multi-label subcellular localization prediction using protein language models. Nucleic Acids Res. 50, W228–W234 (2022).

48. Teufel, F. et al. SignalP 6.0 predicts all five types of signal peptides using protein language models. Nat. Biotechnol. 40, 1023–1025 (2022).

49. Detlefsen, N. S., Hauberg, S. & Boomsma, W. Learning meaningful representations of protein sequences. Nat. Commun. 13, 1914 (2022).

50. Rives, A. et al. Biological structure and function emerge from scaling unsupervised learning to 250 million protein sequences. Proc. Natl. Acad. Sci. U. S. A. 118, (2021).

51. Elnaggar, A. et al. ProtTrans: Toward Understanding the Language of Life Through Self-Supervised Learning. IEEE Trans. Pattern Anal. Mach. Intell. 44, 7112–7127 (2022).

52. Yang, K. K., Fusi, N. & Lu, A. X. Convolutions are competitive with transformers for protein sequence pretraining. Cell Syst 15, 286–294.e2 (2024).

53. Tang, Z. & Koo, P. K. Evaluating the representational power of pre-trained DNA language models for regulatory genomics. bioRxiv (2024) doi:10.1101/2024.02.29.582810.

54. Jumper, J. et al. Highly accurate protein structure prediction with AlphaFold. Nature 596, 583–589 (2021).

55. Abramson, J. et al. Accurate structure prediction of biomolecular interactions with AlphaFold 3. Nature 630, 493–500 (2024).

56. Lin, Z. et al. Language models of protein sequences at the scale of evolution enable accurate structure prediction. bioRxiv 2022.07.20.500902 (2022) doi:10.1101/2022.07.20.500902.

57. Liao, S. E., Sudarshan, M. & Regev, O. Deciphering RNA splicing logic with interpretable machine learning. Proc. Natl. Acad. Sci. U. S. A. 120, e2221165120 (2023).

58. Faure, A. J., Schmiedel, J. M., Baeza-Centurion, P. & Lehner, B. DiMSum: an error model and pipeline for analyzing deep mutational scanning data and diagnosing common experimental pathologies. Genome Biol. 21, 207 (2020).

59. Koo, P. K. & Ploenzke, M. Improving representations of genomic sequence motifs in convolutional networks with exponential activations. Nat Mach Intell 3, 258–266 (2021).

