## Supplementary figures for "Massive experimental quantification of amyloid nucleation allows interpretable deep learning of protein aggregation"

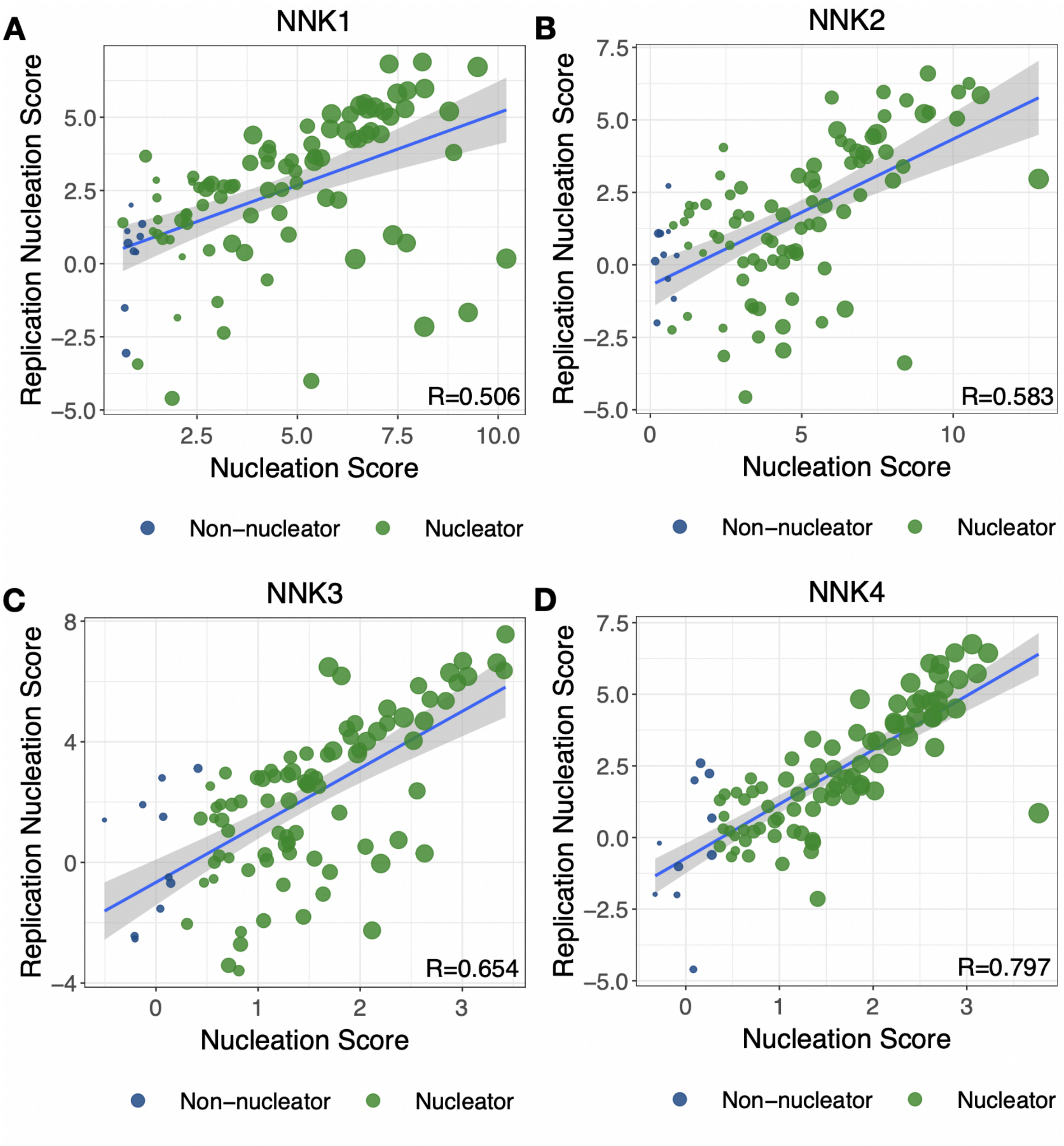


**Supplementary Figure 1 Nucleation scores are replicable across experiments**. Nucleation scores from the corresponding original experiment are plotted on the x-axis, and the replication nucleation score is plotted on the y-axis. Sizes are proportional to the inverse error measurement from the original experiment as reported by DiMSum. The x-axis scores were all calculated independently within their respective experiment—(A) NNK1, (B) NNK2, (C) NNK3, (D) NNK4, the validation set—and altogether in the replication set (n=100), as 100 sequences were taken from each experiment.

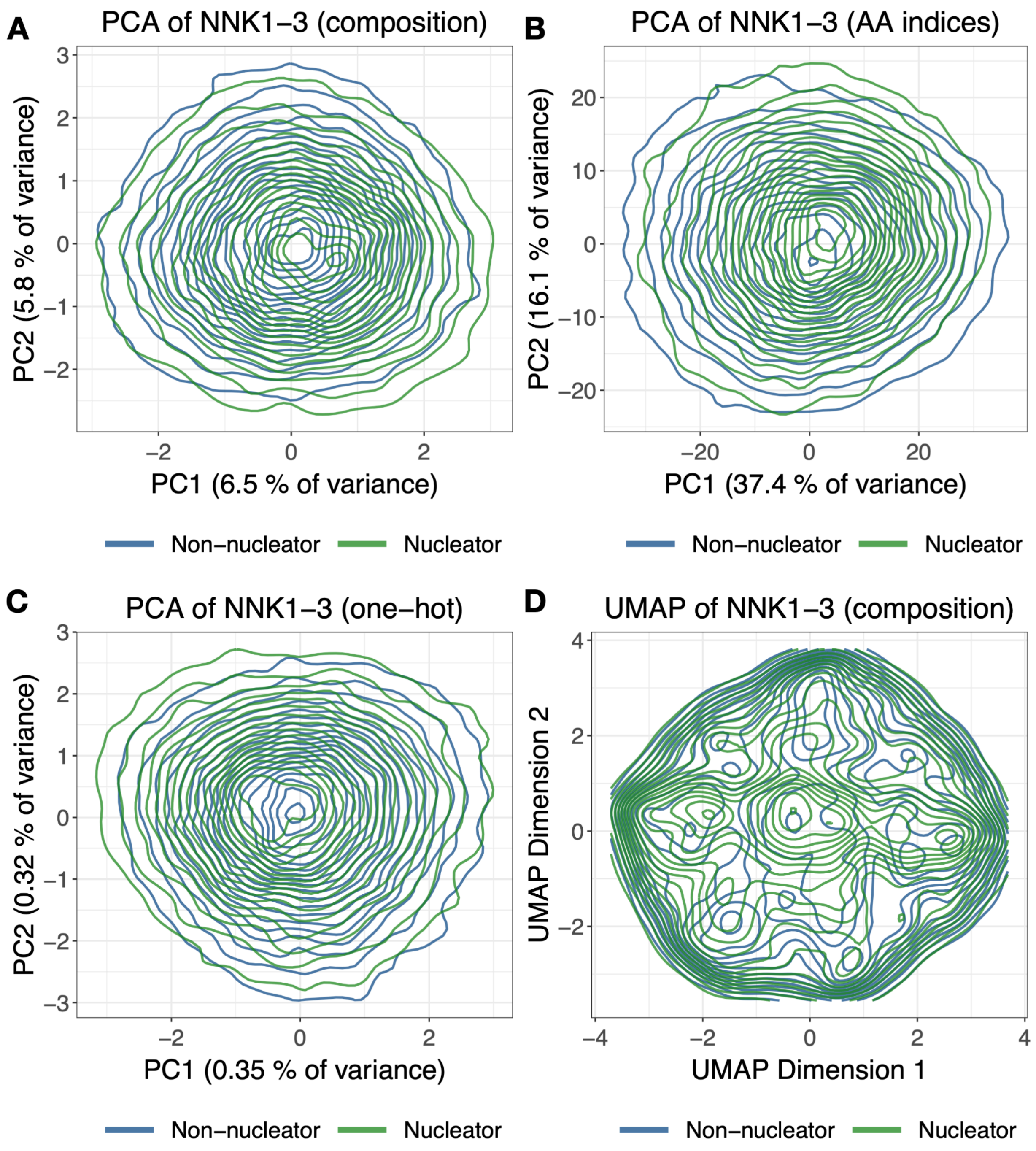


**Supplementary Figure 2 Dimensionality techniques fail to distinguish nucleation status**. (A-C) Contour plots of Principal Component Analysis (PCA) scores on PCs 1 and 2 when sequences are represented by (A) overall amino acid composition, (B) 533 amino acid indices calculated from python package protlearn (C) one-hot (position-maintained) amino acid composition. (D) UMAP projection when using amino acid composition as input.

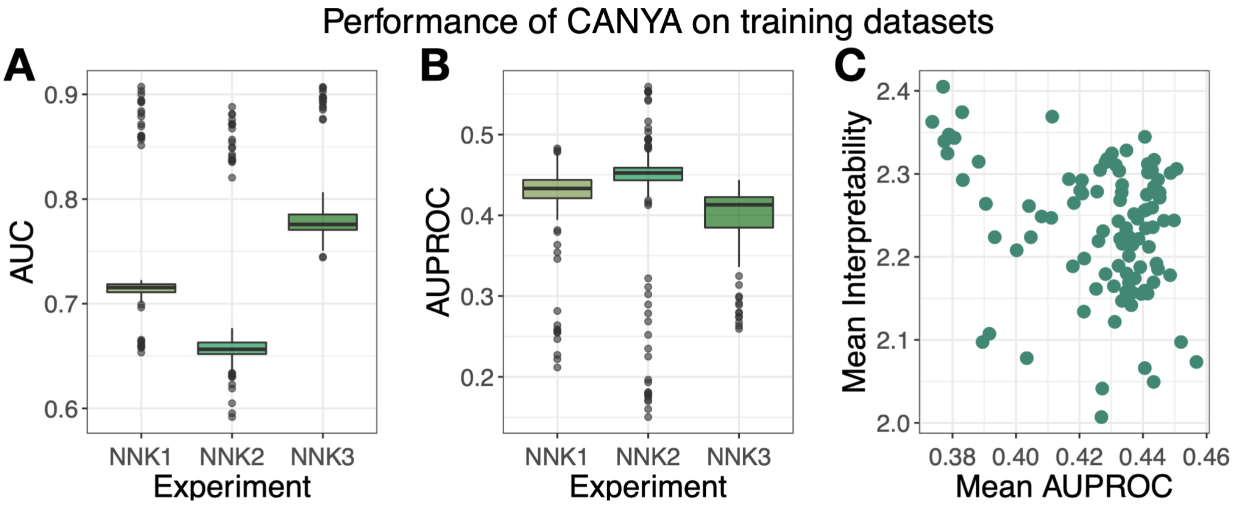


**Supplementary Figure 3 CANYA performance on the held-out test portion of the training datasets**. Evaluation metrics across the all 100 model fits of CANYA. (B) The area under receiver operating characteristic curve (AUC) for held-out testing sequences. (C) The area under precision recall curve (AUPROC) for held-out testing sequences. (D) The interpretability score (KL divergence; Methods) calculated on all held-out test sequences plotted against the mean AUPROC across experiments.

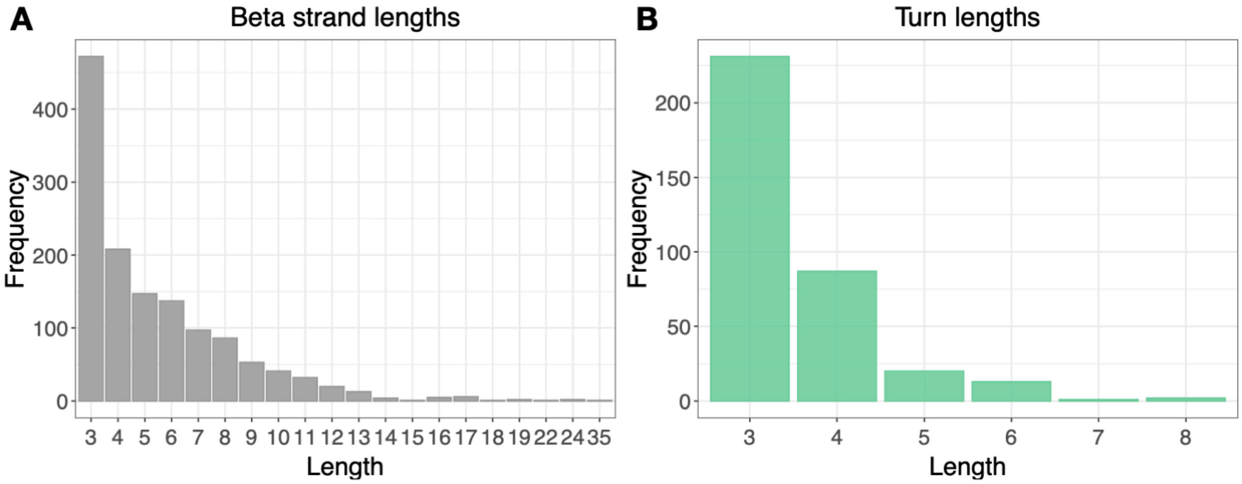


**Supplementary Figure 4 Secondary structure lengths of resolved amyloids**. We downloaded data from the WALTZ-DB data portal, filtered for amyloids, then manually parsed the Uniprot entries of each sequence to obtain the distribution of (A) beta strands and (B) turns across 80 sequences.

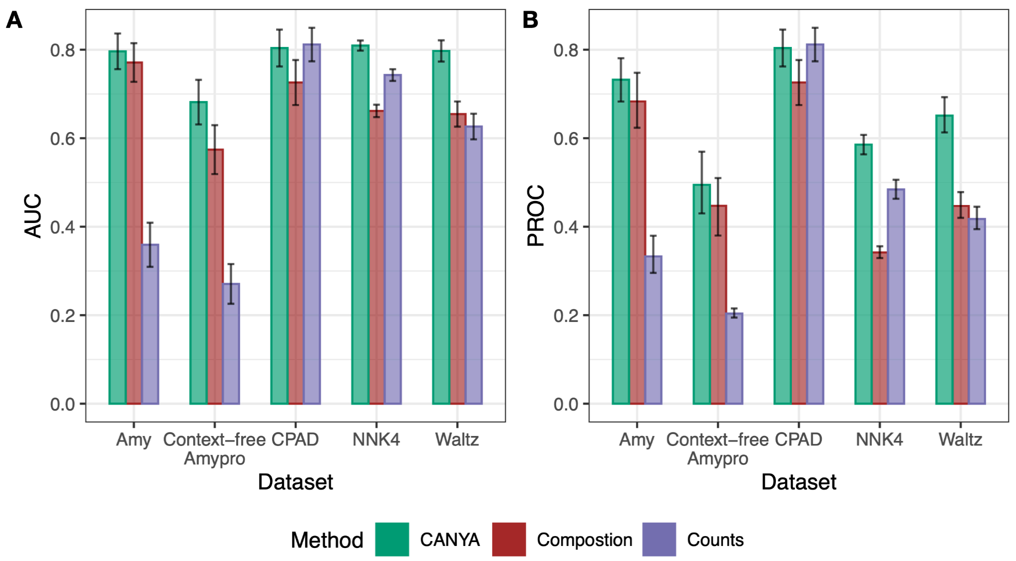


**Supplementary Figure 5 The performance of CANYA compared to simpler models.** Area under the receiver-operating characteristic and precision-recall curves (AUC, AUPROC respectively) of each method on a corresponding testing set (Methods, Supplementary Table 6). "Composition" corresponds to training a simple logistic regression model using amino acid composition (between 0.0 and 1.0, proportions) over the training NNK dataset, and "Counts" corresponds to training the same model with raw, unnormalized amino acid counts (between 0 and 20, integer values).

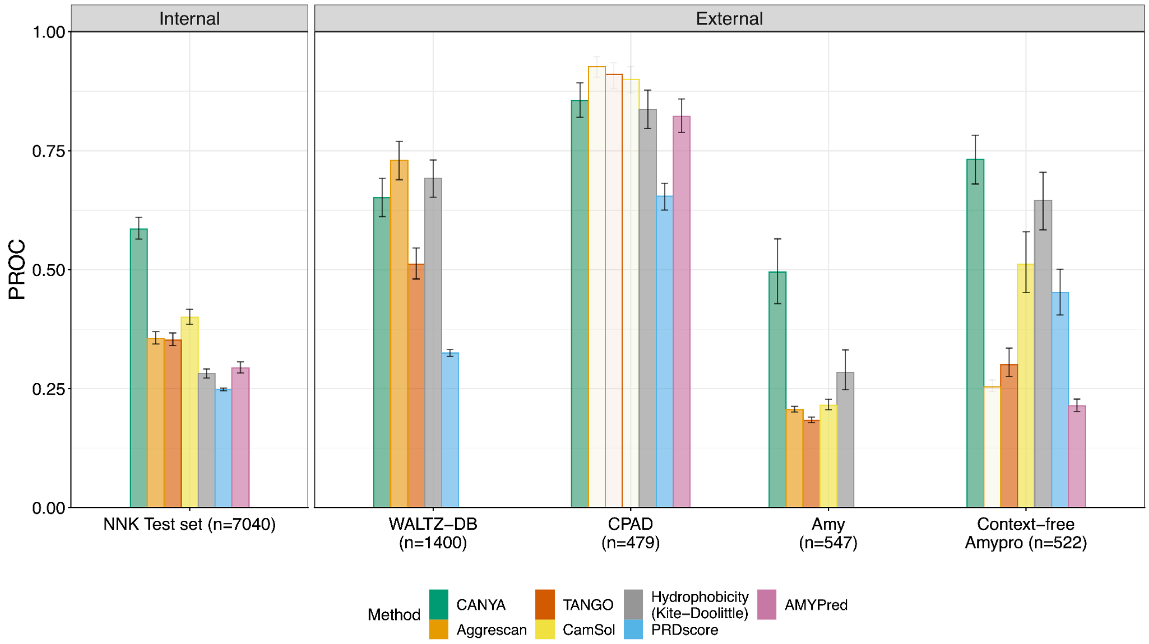


**Supplementary Figure 6 The performance of CANYA compared to previous approaches on testing datasets.** Area under the precision-recall curve (AUPROC) of each method on a corresponding testing set. Low-opacity bars represent cases in which the method used data from the testing dataset to do its training, and thus are not valid out-of-sample evaluations. See text for additional descriptions of datasets (Methods, Supplementary Table 6).

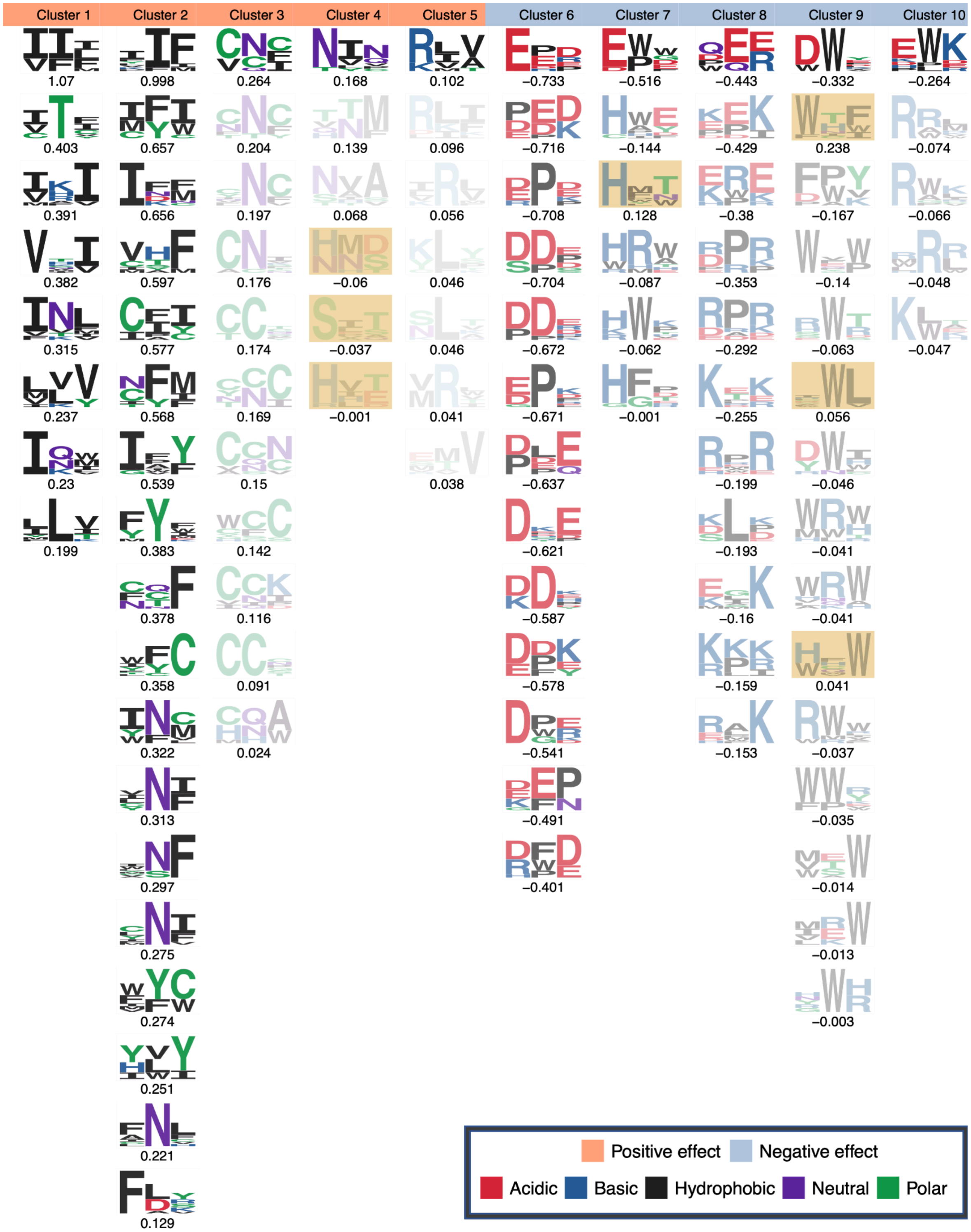


**Supplementary Figure 7 Physicochemical motifs discovered by CANYA prior to performing quality control**. We removed filters from clusters if their GIA effect direction was opposite the sign of the effect of the strongest filter. We highlight which motifs were excluded from downstream xAI analysis in yellow.

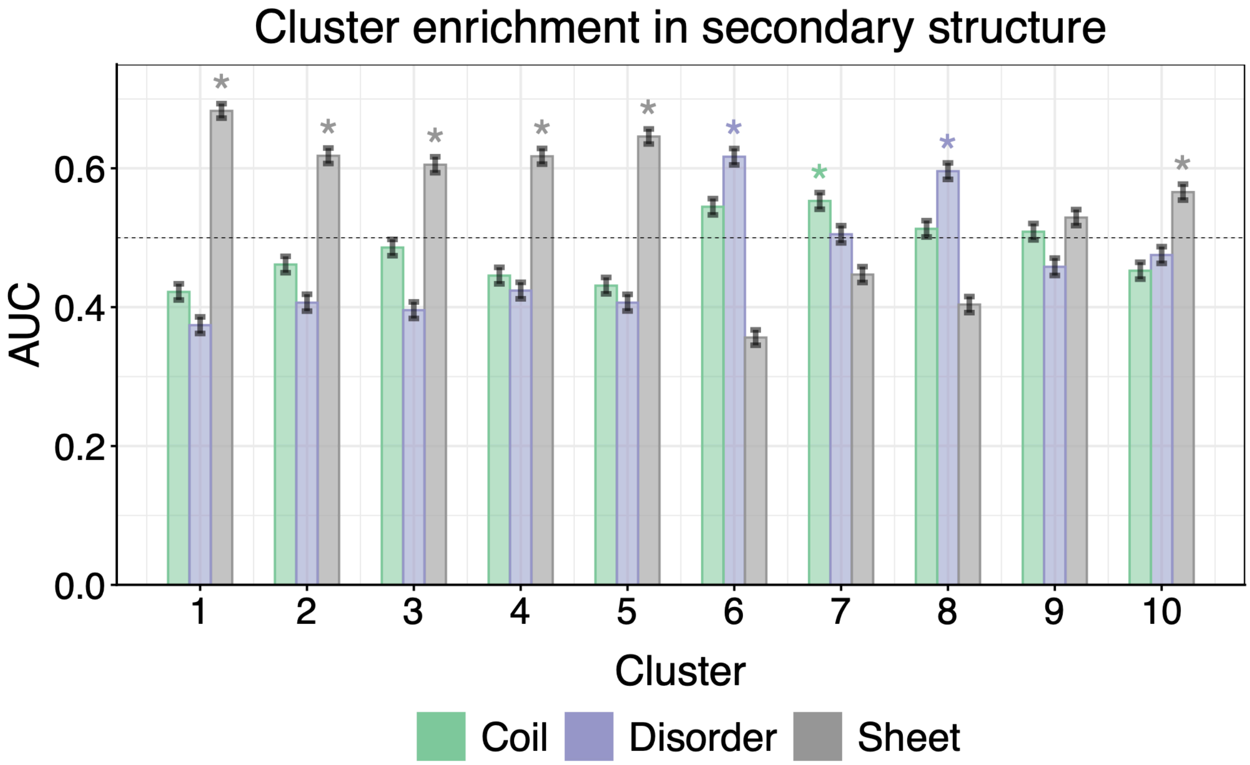


**Supplementary Figure 8 Secondary structure enrichment of motifs discovered by CANYA**. We collected sequences from the StAmP dataset then collected their convolution layer activation energies from CANYA. Across all sequences, we examined whether a specific cluster had higher activation (pattern matching) within a specific secondary structure by calculating the AUC between the activation energy on a specific secondary structure (Methods). Asterisks represent structures for which the enrichment was significantly higher than both 0.50 and the second most-enriched structure.

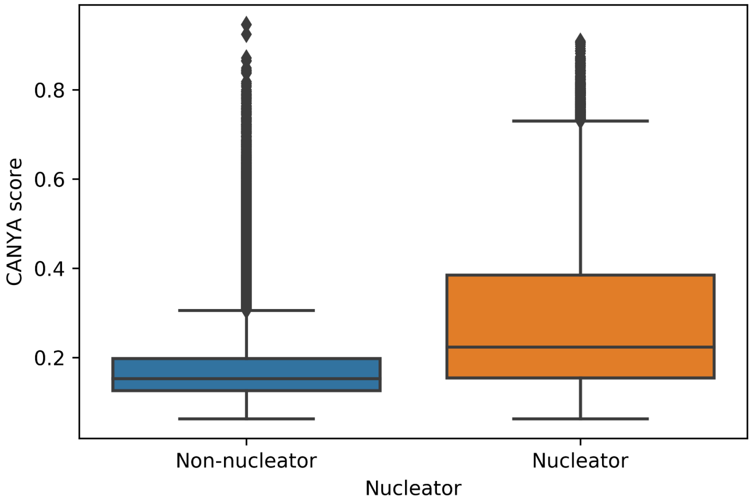


**Supplementary Figure 9 Distribution of CANYA scores across training sequences**. The distribution of output scores for non-nucleators (n=79,910) and nucleators (n=20,826).
